# Understanding the impact of blast-induced traumatic brain injury on brain cellular differentiation and mechanics at nanoscale

**DOI:** 10.1101/2025.04.25.650643

**Authors:** Nabila Masud, Catherine Fonder, Bridget McGovern, Md Hasibul Hasan Hasib, William J. Jackson, Dulce C. Resendiz, Carley River, Sarah A. Bentil, Donald S. Sakaguchi, Anwesha Sarkar

## Abstract

Blast-induced traumatic brain injury (bTBI) causes significant disruptions in cellular and subcellular structures within the central nervous system (CNS) following the application of an extremely large force. The corresponding changes in biomechanical properties and cellular functionalities of neuronal and glial cells (i.e. astrocytes, oligodendrocytes) due to bTBI remain largely unexplored. In this work, controlled shockwave exposure was applied to adult hippocampal progenitor cells (AHPCs) and resultant alterations in nanomechanical, viscoelastic properties, cellular survival, proliferation, and differentiation were examined. bTBI was induced using a custom-designed compression-driven shock tube, and cellular responses to single (overpressure magnitude: 14.5 psi or 100 kPa) and double shockwave exposures (overpressure magnitude: 29.0 psi or 200 kPa) applied in two different directions (from top-to-bottom and bottom-to-top) were analyzed using atomic force microscopy (AFM) and immunocytochemistry (ICC). We observed noticeable changes in cellular mechanics, especially when exposed to a double shockwave from bottom-to-top direction. These variations were characterized by a significant reduction in Young’s modulus, surface roughness, and viscosity. Shockwave exposure from bottom to top direction caused pronounced actin cytoskeletal disruptions compared to top-to-bottom direction. However, ICC results showed that cell viability remained high for both cases of shockwave exposures despite mechanical changes; although the population of oligodendrocytes and immature neurons displayed significant decreases after double shockwave exposure. These findings emphasize the interplay between cellular behavior, resilience of neuronal and glial cells, and cellular nanomechanics in bTBI aftermath, leading to the development of novel therapeutic approaches against bTBI.

## 1 Introduction

Blast-induced traumatic brain injury (bTBI) occurs when extreme mechanical forces, such as those from an explosion, are applied to the brain, causing significant disruption to cellular mechanics at the nanoscale [1, 2]. Unlike blunt force trauma and neurodegenerative diseases, bTBI is specifically caused by sudden and intense forces applied directly to the brain [3–6]. bTBI is a major cause of death and disability in the U.S. and is often caused by explosions in war zones where many service members are injured [7–9]. According to the Glasgow Coma Scale (GCS) as a primary tool, traumatic brain injury (TBI) is commonly categorized by clinical severity. GCS scores classify TBI levels as mild (13-15), moderate (9-12), or severe (≤ 8) [7, 8]. Most bTBIs are mild (concussions) and can be hard to diagnose [10]. Mild TBI (mTBI) can lead to a range of symptoms, such as trouble focusing, blurred vision, irritability, headaches, sleep problems, and depression. In some cases, individuals may develop long-term conditions like post-traumatic stress disorder (PTSD) and chronic traumatic encephalopathy (CTE) [11, 12]. Unfortunately, a lack of understanding of the pathobiological mechanisms underlying TBIs has led to inadequate treatment options [13]. TBIs, including damage from blast injuries, impact the central nervous system (CNS) (see **Fig**.1), which comprises the brain and spinal cord. The neural progenitor cells of the CNS can differentiate into three main lineages: neurons, oligodendrocytes, and astrocytes [14, 15]. Adult hippocampal progenitor cells (AHPCs) are multipotent neural cells in the adult brain, capable of proliferation and differentiation into these three cell types, making them suitable and relevant candidates for the shockwave experiments [14, 16]. Primary bTBI is typically characterized by concealed internal injuries, which makes detecting and assessing the severity challenging. The seriousness of primary bTBI is influenced by factors such as the distance from the explosion, its intensity, and duration [17]. Damage results from an overpressure wave passing through tissues, and the CNS is particularly vulnerable. Therefore, gaining a fundamental understanding is imperative for developing targeted diagnostic and therapeutic strategies to address current clinical limitations in TBI.

**Fig. 1.**
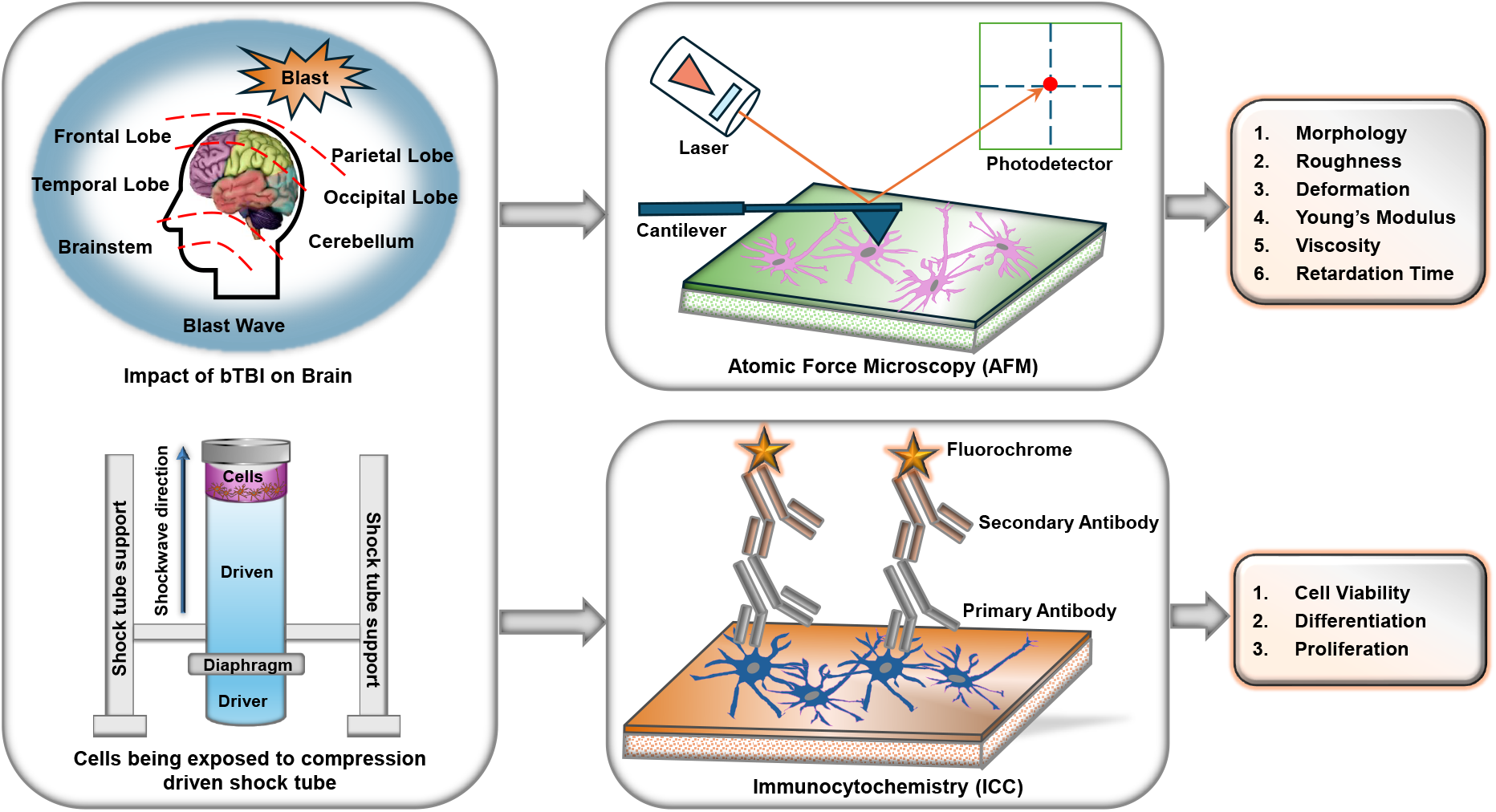
Schematic overview of the study objectives and experimental workflow. The left section illustrates the effects of blast-induced traumatic brain injury (TBI), along with the custom setup used to apply shockwaves (controlled blast pressures) to cells. The second section details the investigation procedures, showing (top right corner) the Atomic Force Microscopy (AFM) setup for analyzing structural, nanomechanical, and viscoelastic properties and (bottom right corner) the immunocytochemistry (ICC) setup for complementary cellular analysis.

The cell type, along with the extracellular matrix, plasma membrane, transmembrane receptors, cytoskeleton, and nuclear architecture, all affect how cells respond to applied forces, enabling neurons and glial cells to react to their physical surroundings. However, when the forces become too strong, like in TBI (see **Fig**.1), these components may respond differently or fail to function, leading to long-term cell damage and neurodegeneration [18, 19]. Previous research has made noteworthy strides in analyzing neuronal plasma membrane disruption across various injury models and revealed a positive correlation between insult severity and the degree of membrane disruption [20]. Recent studies also have demonstrated that mechanical forces can disrupt the cytoskeleton and impair the differentiation of neural progenitor cells, leading to long-term dysfunction. For example, [21] found that stem cells subjected to mechanical trauma experienced significant alterations in their ability to proliferate and differentiate, underscoring the sensitivity of these cells to mechanical stimuli. Furthermore, a study by [22] indicated that mechanical forces play a critical role in modulating stem cell behavior by influencing their proliferation and differentiation. This dynamic regulation has significant implications for regenerative medicine, as understanding these mechanisms can enhance the development of therapeutic strategies. These findings highlight the importance of investigating how mechanical forces affect the nanomechanical properties of stem cells following trauma.

Recent advances in immunocytochemistry (ICC) techniques have enabled researchers to more precisely measure post-trauma cellular responses, particularly in relation to neural regeneration [23]. For example, [24] used ICC to analyze brain specimens from 11 TBI patients and showed an increased expression of neural stem/progenitor markers in the perilesional cortex. This suggests a potential neurogenesis response in the adult human brain after a traumatic brain injury. Additionally, Atomic Force Microscopy (AFM) is a valuable tool for studying mechanical changes in cells. Its ability to provide high-resolution mapping of nanomechanical properties is particularly important for stem cell research, as it enables the measurement of critical factors such as Young’s modulus, deformation, viscosity, and retardation time that are key indicators of cellular health and function [25, 26]. AFM has been extensively used in brain cellular mechanics research to measure nanomechanical properties, providing insight into how trauma affects cellular structure [27]. Magdesian et al.[28] utilized AFM to demonstrate that axonal degeneration resulting from traumatic brain injury and nerve compression leads to significant biomechanical changes in axons like neural damage and dysfunction. These studies highlight the ongoing need for research to unravel the complexities of cellular responses to traumatic injuries, laying the groundwork for more effective diagnostic and treatment interventions [29]. This study incorporated unique parameters, such as subjecting cells *in vitro*, instead of live animal models to the blast wave, and positioning substrates without being attached to a blast setup, as seen in animal studies. AHPCs were subjected to a shockwave generated through an extended tube from a distance, potentially allowing air blast and sound release to affect the cells. These distinctive aspects enabled a more thorough investigation of the neural cells’ responses to the applied shockwave. Thus, we aimed to assess the nanomechanical changes and functional outcomes of AHPCs post-trauma (single and double shockwave exposure) by utilizing AFM and ICC to evaluate their proliferation, differentiation, and viability (see **Fig**.1). This integrated approach enabled us to explore how the nanomechanical properties of AHPCs correlate with functional behaviors following exposures to shockwaves of two different magnitudes and two different directions, contributing to the development of therapeutic strategies aimed at mitigating long-term damage caused by TBI.

## 2 Results and Discussion

### 2.1 Shockwave Generation and Propagation in ‘Top-to-Bottom’ and ‘Bottom-to-Top’ Directions

In our study, a customized compression-driven shock tube was utilized to generate high-pressure shockwaves for exposing AHPCs and investigate the effects of blast waves at the cellular level. The Schedule 80 shock tube had a 3/4-inch inner diameter and was comprised of a 4-inch driver and 30-inch driven section, separated by a diaphragm [30]. This diaphragm served as a barrier between the driver section and the driven section containing the test region with the APHCs. In shockwave-treated cells, the shock tube system operated by filling a driver section with compressed air until a Dead Soft aluminum diaphragm was ruptured, creating a shockwave that propagates from one end of the tube (i.e. driver section) to the other (i.e. driven section) (see **Fig**.1 bottom left). Control cells were retrieved from the incubator at the same time as those subjected to the shockwave to ensure consistent results and accurate comparative analysis.

For the shockwave exposure, there were a total of four conditions: two for the shockwave overpressure magnitudes (single and double) and two for directional exposure (top-to-bottom and bottom-to-top). A single shockwave utilized an overpressure amplitude of 14.5 psi (100 kPa); a double shockwave repeated the exposure twice for 29 psi (200 kPa). The shockwaves were applied from two different directions with respect to the petridish holding the cells. In one case, the pressure was applied from above, and it got transmitted successively through several barriers—the lids of the petridish, the air space above the medium, the cell culture medium, and finally, the cells themselves. In another instance, the application of pressure from beneath directs it straight away at the base of the petridish for a more direct impact on the cells (see **Fig**.2).

**Fig. 2.**
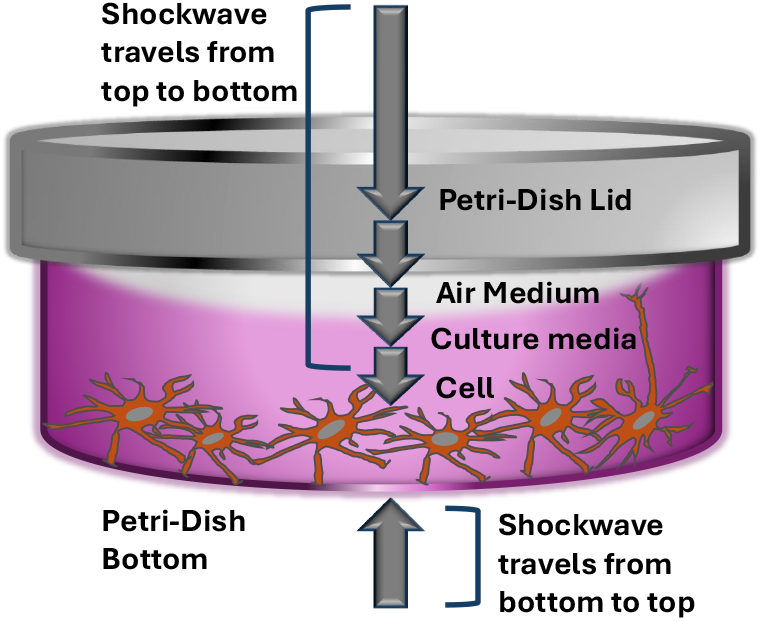
Experimental setup for applying shockwaves to cells in a petridish. Shockwaves was delivered either downward (top-to-bottom) or upward (bottom-to-top).

### 2.2 Morphological Changes in Cells Following Shockwave exposures

Besides directly controlling the motility of cells, signaling, and intracellular transport, the cytoskeleton plays an important role in maintaining cellular structure [31]. Significant changes in cellular morphology could hence occur upon exposure to a shockwave, in which the nature of the changes might differ depending on the environmental load. More interestingly, the structure of the cytoskeleton is especially sensitive to the effects of shockwaves. In our study, first, we investigated the morphological changes of AHPC cells subjected to shockwaves with varied intensities and orientations. With this approach, it was possible to explain how these variable parameters influence the stability of the cytoskeleton and overall cellular morphology. In control cells that were not exposed to shockwaves, the actin cytoskeleton was strongly assembled into actin-rich structures in specific cellular areas. The filaments of actin were assembled into clear assemblies and striations, which contributed much to the mechanical strength, shape-changing ability of the cell, and structural integrity to the cell in order for it to withstand all mechanical stresses (**Fig**.3(A,D)). Stress fibers, known as temporary contractile bundles, formed by actin and myosin, connected focal adhesion sites on the plasma membrane [32]. Such organized cytoskeletal structures are essential for cellular resilience, as they help absorb and distribute external forces, maintaining the cell’s mechanical properties and its ability to adhere to its substrate.

**Fig. 3.**
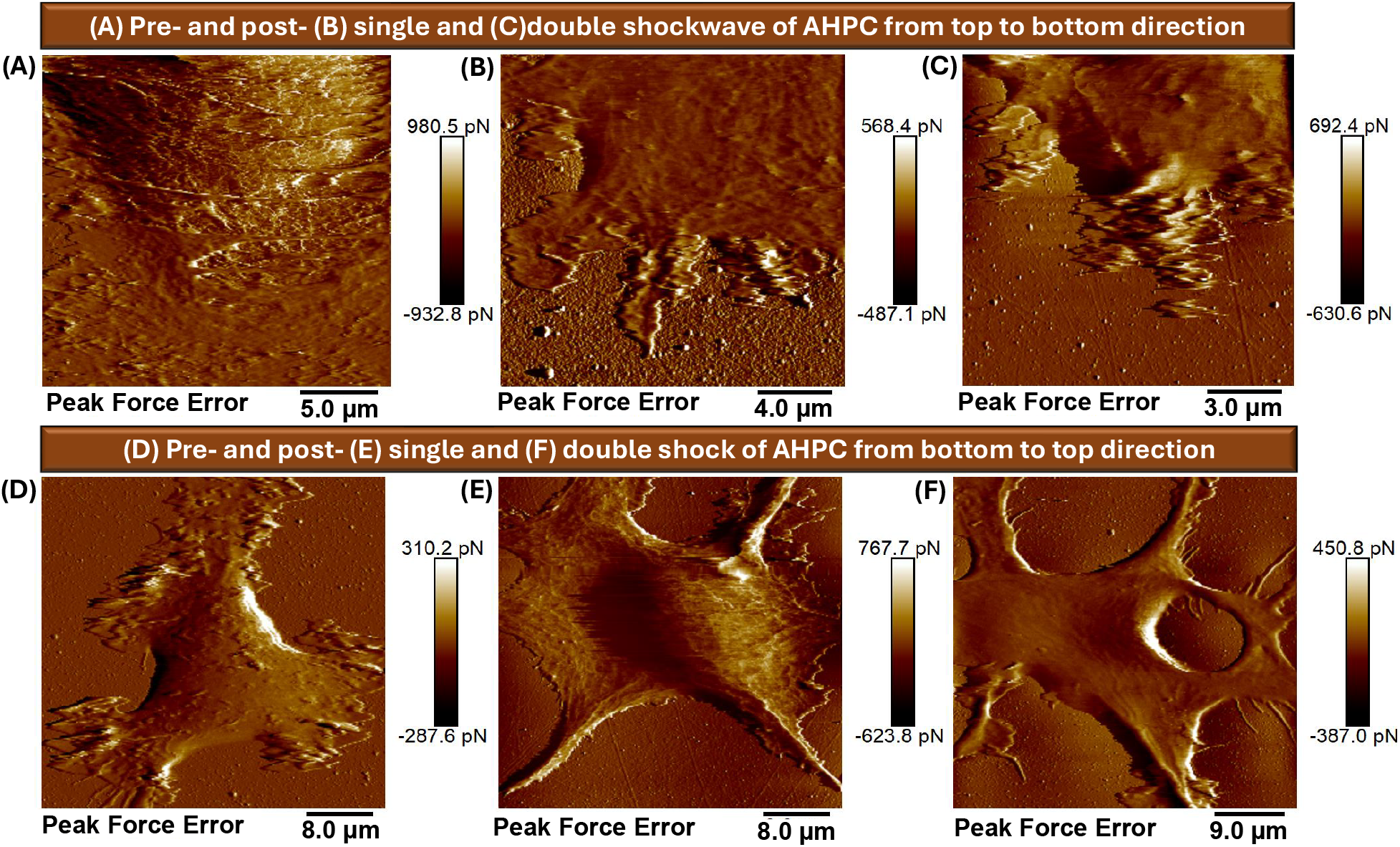
Peak force error images of cells obtained using AFM under different shockwave exposure conditions. The top section shows images of control cells(A) alongside cells exposed to a shockwave from the top at 14.5 psi(B) and 29 psi(C). The bottom section presents control images(D) along with images of cells exposed to a shockwave from the bottom of the cell dish at 14.5 psi(E) and 29 psi(F). This comparison highlights morphological changes in cell surfaces due to directional shockwave exposure at varying pressures.

These cytoskeletal components, initially forming a cohesive and dense network with striations; upon shockwave exposure, started becoming weaker and became aggregated, primarily around the cell periphery. Notably, actin filaments smoothened, and also retracted from the perinuclear area towards the edge of the cell (see **Fig**.3(F), creating an uneven cytoskeletal distribution. The retracted cells showed a rearrangement of their actin filaments into elongated stress fibers in the filopodia [33], likely an adaptive response to structural damage from the shockwave. Moreover, when subjected to shockwaves delivered from the bottom of the cell dish, the cells exhibited globular morphology and blebbing, even in response to forces less than 100 pN (see Supplementary **Fig**.A8), which are indicative of mechanical stress and cytoskeletal disruption. Over time, these treated cells continued to alter in shape, becoming more globular, and blebs formed on their surfaces. We have also noticed that the cytoskeletal network of cells began to recover, restoring their initial shape approximately 3 hours post-exposure under incubation conditions indicating the general cell health restoration (see Supplementary **Fig**.A8). In this respect, Moosavi-Nejad et al. [34], documented that the exposure of human renal carcinoma cells to shockwave results in disorganization of the cytoskeleton with an increased amount of reconfiguration in actin and tubulin and practically no change in vimentin filaments. Such structural reorganization, according to these authors, may be due to the effects of localized cavitation where high-intensity forces would be produced at a very focal cellular site. Although our shockwave exposure differed by being applied through the cell culture dish rather than directly at cellular sites, similar cytoskeletal alterations were observed in our study. This suggests that even shockwaves of moderate magnitudes (single shockwave of 14.5 psi or 100 kPa magnitude, double shockwave of 29.0 psi or 200 kPa magnitude) can induce structural modifications across the cytoskeleton, potentially due to distributed pressure across the cell’s attachment points on the culture substrate.

In a further study of the effect of shockwave exposure on cell surface conditions, our analysis of the cellular surface roughness showed that shockwave exposure reduced the cell membrane roughness with increases in both pressure and time. Indeed, a top-to-bottom shockwave of 14.5 psi magnitude applied for 1.76 ms duration did not have a significant reduction in surface roughness with an RMS of 11.17 ± 1.17 nm compared to the control cells, which had an RMS value of 11.69 ± 1.69 nm. However, increasing the pressure of the shockwave magnitude to 29 psi with a duration of consecutive twice exposure of 1.76 ms produced noticeable smoothing, with an RMS roughness measured at 9.21 ± 1.44 nm, as displayed in **Fig**.4(A)). Even more interestingly, bottom-to-top propagating shockwave produced further smoothing and weakening of the cytoskeletal network, where single and double shock exposures reduced cell surface roughness to 9.83 ± 1.10 nm and 8.14 ± 1.08 nm respectively, as shown in **Fig**.4(B)). The obtained results, therefore, form the ground of proof that the modification of surface roughness after bTBI is directly connected with the resulting cytoskeletal damage; hence, shockwave-induced deformation influences cell membrane characteristics along with cytoskeletal structure.

**Fig. 4.**
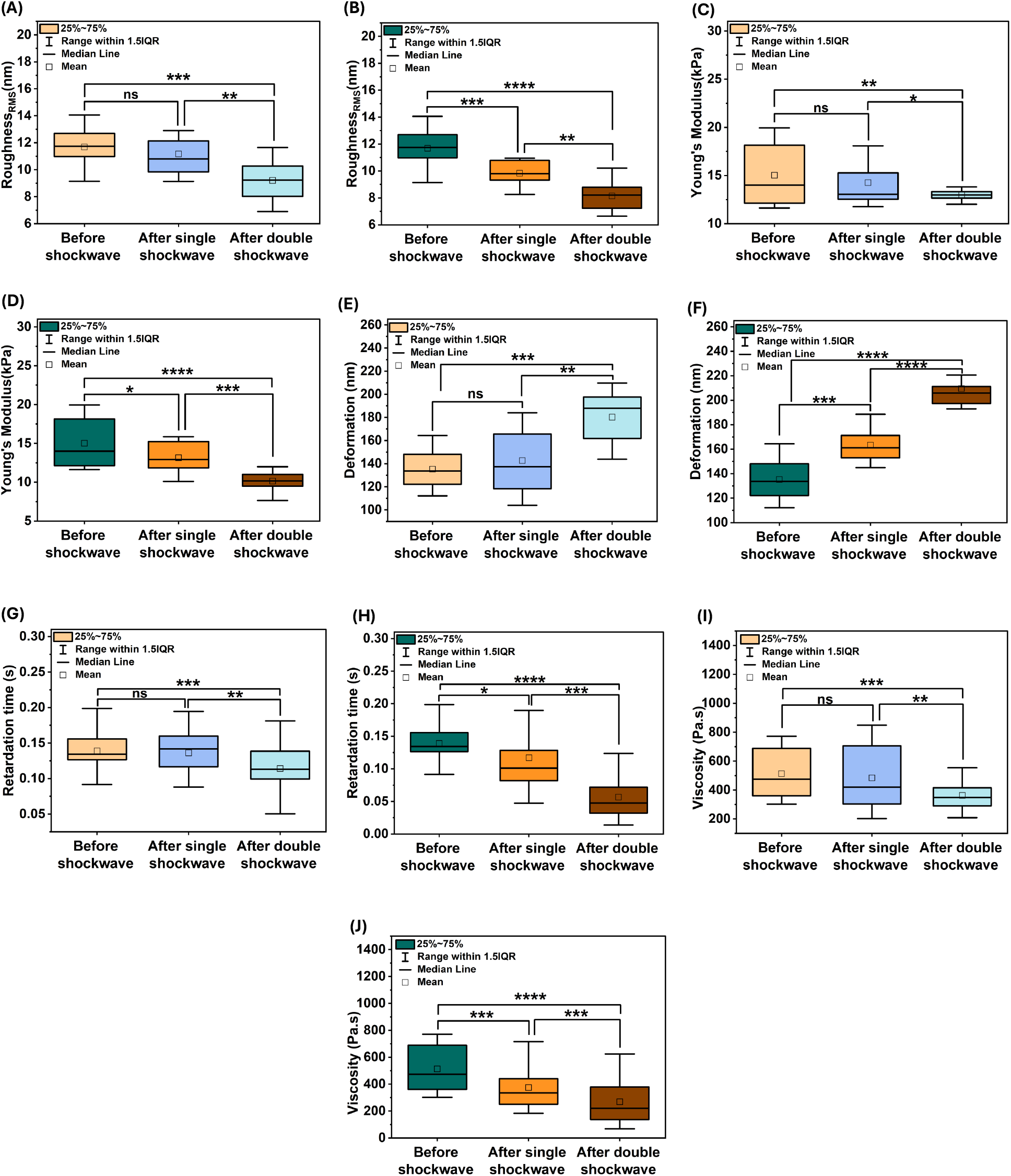
Effects of shockwave exposure from top-to-bottom and bottom-to-top directions on the morphological and mechanical properties of AHPCs. Box plots illustrate comparisons across three conditions—before shockwave, after single shockwave, and after double shockwave exposure—under top-to-bottom shockwave exposure for (A) surface roughness, (C) Young’s modulus, (E) deformation, (G) retardation time, and (I) viscosity. Similarly, the impacts of single and double shockwave exposure from the bottom-to-top direction on AHPCs are shown in box plots for (B) surface roughness, (D) Young’s modulus, (F) deformation, (H) retardation time, and (J) viscosity. Significant decreases in roughness, Young’s modulus, viscosity, and retardation time, along with an increase in deformation, are observed after shockwave exposure, particularly following double shockwave exposure in both cases. “X-X” represents the upper and lower extremes of the data, while the error bars extend to a range of 1.5 times the interquartile range (IQR). The mean is depicted by the small box, and the median is indicated by the black line. Statistical significance is denoted as follows: ns (not significant), * *p <* 0.05, ** *p <* 0.01, *** *p <* 0.001, **** *p <* 0.0001.

Previous *in vivo* studies have demonstrated that the extracellular glutamate level increases following TBI [35, 36]. This excitotoxicity mediated by glutamate significantly contributes to altered neuronal cytoskeletal stability subsequent to injury. Such changes are often attributed to the calcium influx initiation by glutamate excitotoxicity [37, 38]. Thus, increased intracellular calcium has been linked to decreased extracellular levels of Ca^2+^ following even mild TBI [37]. Activated via calcium flux, the calpains specifically degrade cytoskeletal proteins and lead to behavioral impairments [39, 40]. Further, these changes lead to depolymerization of MTs and NFs [41], which correspond to decreased levels of MAP2 and NFs, thus contributing to cytoskeletal instability and cellular morphology changes [42]. Combined with the findings of other TBI literature, our findings here indicate that the cytoskeletal damage and reduction in surface roughness resulting from bTBI stem from a direct mechanical disruption. This could be a subsequent biochemical cascade, such as glutamate-induced excitotoxicity and calcium influx. The results also emphasize the multifaceted effects of shockwaves on cellular morphology, with interdependent mechanical and biochemical processes compromising the integrity of the cytoskeleton and overall cellular architecture following bTBI.

### 2.3 Influence of Shockwave Application on AHPC Cell Mechanics

TBI is a complex interaction of external physical forces and unique biomechanical properties of the brain. It, therefore, becomes important to understand the mechanical response of the brain to these external loads. Some of the basic concepts that define the mechanical properties of tissues or cells include stress, strain, and Young’s modulus. The force applied over an area engenders strain, the ratio of post-force dimensions to the original shape, or the deformation ratio of an object [43, 44]. Brain tissue is a viscoelastic material, and it exhibits stiffness - or softness - based on the speed at which a force is applied. Faster loading leads to higher stiffness, whereas slow loading tends to produce softer, fluid-like behavior [45]. The unique viscoelastic nature of brain tissue underlies the response of brain tissue to various loading scenarios associated with bTBI and points out its dual nature: elastic under rapid stress and viscous under prolonged stress. Thus, in this respect, the elastic and viscous properties of the AHPC, as well as the deformation and retardation time, were investigated in this paper for a better understanding of their contribution to the mechanical responses of AHPC under various injury conditions.

#### 2.3.1 Elastic properties

Injuries can dramatically alter the mechanical properties of tissue. For instance, one study that explored cortical impacts in mice learned that the shear stiffness of the injured area of the brain was significantly reduced at the time of injury and remained so for days [46]. This change in stiffness suggests that the mechanical integrity of brain tissue is compromised following traumatic events. Therefore, to investigate the effects of cellular mechanics, we varied the mechanical stresses applied to cells by adjusting both the pressure and the duration of exposure. In our experiments, cells were exposed to shockwaves of two different magnitudes, two different directions, and two duration. We also performed control experiments with cell petridishes that were not exposed to shock waves. Immediately after shockwave exposure, the cell dishes were then transferred onto the AFM stage to capture detailed nanomechanical properties. Fitting of the force-distance curves with the Hertz model **(Eq.(4))** was applied to obtain the changes in Young’s modulus.

Our result indicated that, after the shockwave exposure, there was a significant decrease in Young’s modulus, meaning mechanically, the cell was softer than that before the shockwave. For control cells, the resting Young’s modulus of the cells was 15.023 ± 3.17 kPa. After a single shockwave exposure, this value decreased to 14.315 ± 2.8 kPa, and further dropped to 12.984 ± 1.2 kPa after double shockwave exposure (see **Fig**.4(C)). The reduction in stiffness after a single shockwave was insignificant, whereas the decrease in stiffness after double shockwave exposure was significant, showing that prominent mechanical changes occur at the cellular level. Additionally, we analyzed cell deformation after shockwave exposure, which showed an increasing trend compared to the control. The average deformation for control cells was 135.23 ± 17.20 nm, which increased to 142.65 ± 27.34 nm after single shockwave exposure and to 180.18 ± 21.14 nm after double shockwave exposure (see **Fig**.4(E)).

In order to study the direct mechanical effect of shockwaves on adherent AHPCs, we modified the experimental configuration and placed the shock tube under the cell dish. This kind of configuration allowed for a more direct interaction of the shock wave with the cell layer attached at the bottom of the polystyrene dish, where the shockwave had to pass through just one polystyrene layer. Under these conditions, single shockwave exposure was found to be sufficient to cause a detectable change in cell stiffness, with a Young’s modulus of 13.14 ± 3.16 kPa. When the same configuration was exposed to a double shockwave, Young’s modulus further decreased to 10.152 ± 1.12 kPa, as depicted in **Fig**.4(D). The result of deformations in this configuration was similarly reflected when single shockwave exposure yielded an average deformation of 163.454 ± 14.17 nm and double shockwave exposure yielded 209.483 ± 19.32 nm, as shown in **Fig**.4(F). These results, therefore, indicate that the mechanical properties of AHPCs were significantly affected by double shockwave exposure from top to bottom direction and single and double shockwave exposure from bottom to top directions, leading to further cell softening and deformation.

#### 2.3.2 Viscous properties

The AFM Ramp and Hold file is one of several approaches aimed at studying the viscoelastic properties of cellular surfaces by assessing the dynamic response of a cell to an applied periodic strain by monitoring the loading force. Wu et al. [47] illustrated the use of AFM in creep measurements, where the base displacement of the cantilever is controlled to maintain a constant loading force while observing the displacement over time. Several other studies have extended this knowledge. As an example, a stress relaxation study was carried out by Darling et al. [48] on zonal articular chondrocytes by keeping the base of the cantilever fixed and monitoring the shift in loading force with respect to time. With these advances, little is yet understood regarding how changes in loading rate affect the process of relaxation and how this relates to the elastic properties of cells, especially in the bTBI aftermath. Therefore, in our study, we began to investigate the relaxation behavior of AHPC cell surfaces under different bTBI conditions through measurements of the relaxation response at varied scan rates and loading velocities. By varying the loading rate, we aimed to observe how exposure to shockwaves influences the cellular behavior in the relaxation process. The schematic of the stress relaxation measurement setup using AFM is shown in **Fig**.5 for the control and shockwave-exposed cases. The AFM tip was approached towards the cell surface at scan rates of 1 Hz, 2.03 Hz, 3.05 Hz, 4.07 Hz, and 5.09 Hz, with corresponding loading velocities of 10 *µ*m/s, 20.3 *µ*m/s, 30.5 *µ*m/s, 40.7 *µ*m/s, and 50.9 *µ*m/s, while maintaining a constant cantilever base displacement for 2 seconds. The tip was then retracted at the same rate as the approach.

**Fig. 5.**
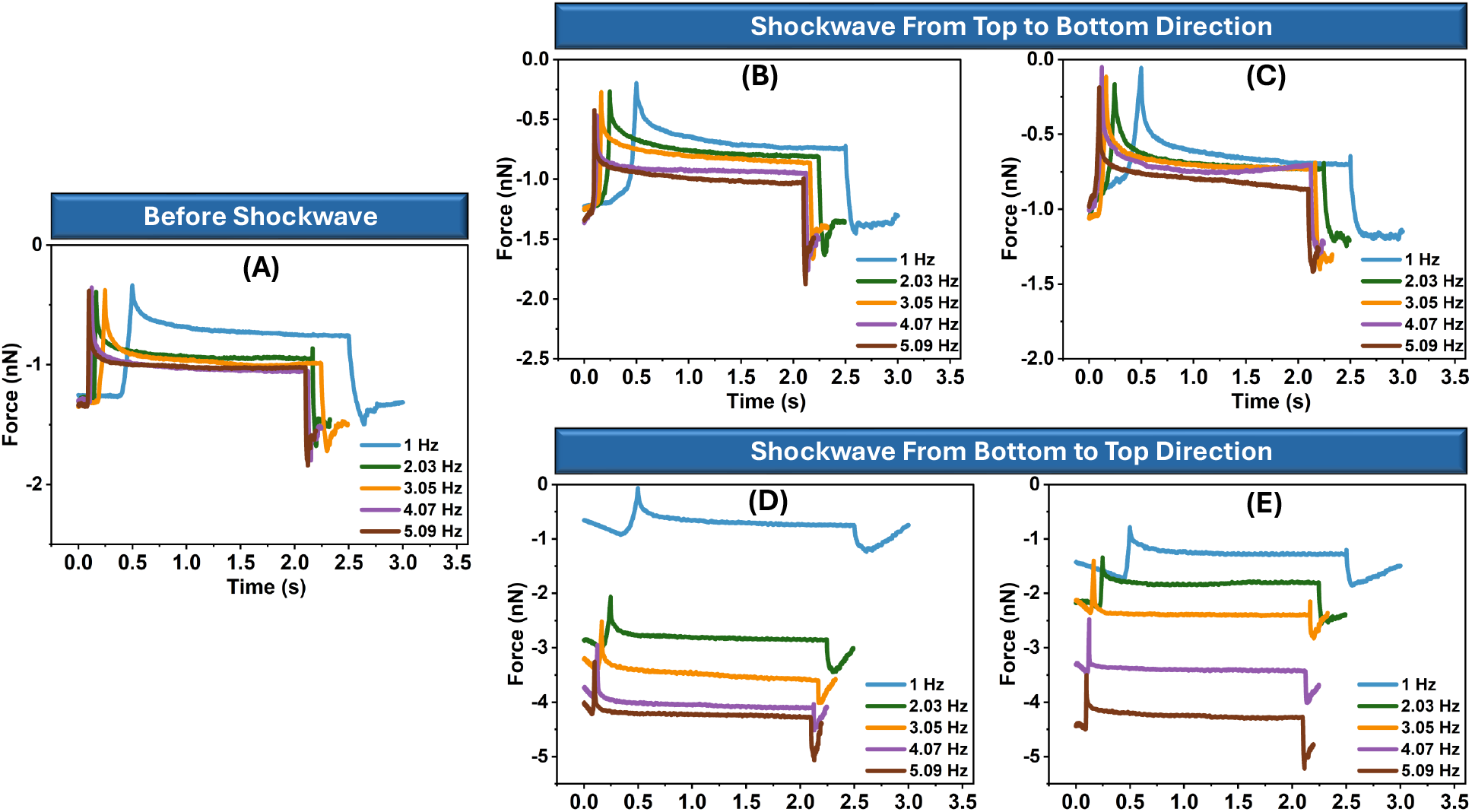
Representation of force-time curve measured in AHPCs before shockwave exposure (A), after a single shockwave (B, D), and after double shockwave exposures (C, E) from both top-to-bottom (B, C) and bottom-to-top (D, E) directions, with the scan rate ranging from 1 Hz to 5.09 Hz. The tip approached and detected the cell surface, and during the 2-second holding period, the loading force began to decrease because of the cell’s viscoelastic properties.

Deflection signals of time series captured during the experiment are shown in **Fig**.5, which plots the force vs. time curve for AHPCs during the stress relaxation experiment. It was observed that the AFM tip approached the cell until the loading force reached a threshold set point, showing afterward a nonlinear decrease in the loading force due to the viscoelastic properties of the cell. This slow decay was more pronounced at the low loading rate. It was also observable that with increasing scan rate in each case, the cells behaved stiffer. For example, by observing the approach curve in **Fig**.5(A-E), at a 5.09 Hz scan rate, cells were showing more stiffness compared to that at 1 Hz scan rate. The cell did not have enough time for viscoelastic relaxation because of the fast movement of the probe. Thus, the stress relaxation graph depicted a much steeper and shorter decay phase due to the cell’s response to high scan rates being much faster and more elastic. On the contrary, when the scan rate was lower, the cell had more time to respond to the applied stress showcasing a slower, more extended relaxation curve. As a consequence of this slow decay, a greater amount of the viscous properties of the cell were captured, thereby giving a more gradual drop in the relaxation curve. Following the 2 sec holding period, the AFM tip was retracted, resulting in force curves that separated at the maximum indentation point. Compared to the control behavior, the relaxation process of AHPCs was similar across all cases, whether the shockwave was applied from top-to-bottom or bottom-to-top, regardless of the exposure duration of 1.76 ms or consecutive two exposures of 1.76 ms each. In all conditions, an increase in the loading rate correlated with an increase in the applied force, indicating that the viscoelastic properties of the cells were consistently responsive to mechanical stress. This suggests that the inherent cellular mechanics were still preserved despite the directional or temporal variations of shockwave exposure, and the relaxation process took place within a time frame of less than 2 seconds.

To quantitatively assess the relaxation behavior, we employed the Kohlrausch-Williams-Watts (KWW) function **(Eq.5)**, an empirical model used to describe the dispersion processes in viscoelastic systems. The relaxation time (*τ*) for control cells was determined to be 0.139 ± 0.029 s. Following a single shockwave exposure (top-to-bottom), the relaxation time showed minimal change at 0.136 ± 0.025 s, whereas double shock exposure resulted in a decrease to 0.114 ± 0.028 s (see **Fig**.4 (G)). In contrast, the bottom-to-top shockwave exposure produced more significant effects: single shockwave measurements yielded a relaxation time of 0.117 ± 0.088 s, and double shockwave reduced those times even more to 0.056 ± 0.034 s (**Fig**.4(H)) respectively. The analysis of cellular viscosity also revealed notable trends. Control cells exhibited a viscosity of 512.98 ± 163.95 Pa · s. After a top-to-bottom single shockwave exposure, the viscosity decreased to 484.49 ± 209.75 Pa · s, with a further reduction to 362.46 ± 95.63 Pa · s respectively following double shock exposure (**Fig**.4 (I)). For cells exposed to bottom-to-top shocks, the viscosity following a single shock was 374.22 ± 149.01 Pa · s, and this value dropped significantly to 268.61 ± 164.33 Pa · s respectively after a double shock (**Fig**.4(J)). These findings indicate a significant impact of both the direction and intensity of shockwave exposure on the viscoelastic properties of AHPCs. The decrease in relaxation time and viscosity suggests that double shock exposures, regardless of direction, induce substantial softening and adaptive viscoelastic responses in the cells. The results highlight that top-to-bottom shockwaves exhibit a gradual decrease in viscoelastic parameters, while bottom-to-top shocks result in a more pronounced reduction.

In all, the mechanical effect of TBI at the cellular level encompasses changes in the neuronal cytoskeleton and ECM. The neuronal cytoskeleton is remodeled in response to TBI, especially by the imbalance of neurotransmitters such as glutamate, which, is released in response to potassium flux and neuronal depolarization following injury [49]. This glutamate-evoked remodeling impinges on the organization of microtubules and, by inference, dendritic growth and retraction, as seen in hippocampal neurons. These ion channels, including the NMDA receptors, further interact with the cytoskeleton and membrane lipids, exhibiting mechanosensitivity either directly or indirectly through conformational changes brought about by force itself [50, 51]. Due to the mechanosensitivity, neurons and glial cells are capable of dynamic response to stress, an important aspect in preventing damage under traumatic forces. Ion channels, for example, are mechanically modulated and allow the regulation of cellular responses to stress and pressure that may influence changes to the plasma membrane after TBI [52]. Moreover, the fact that shockwaves propagate through the brain so fast affects cellular and subcellular structures, which modifies mechanical properties and, consequently, a loss of elasticity and misalignment in axons, as confirmed by cortical impact and stretch injury studies [46, 53]. Furthermore, such disruptions in the integrity of the cytoskeleton following TBI have been illustrated on various levels: from changes at the protein level, such as MAP2 and neurofilaments, to alterations in dendritic morphology for some time following injury [54, 55]. Therefore, this altered structure of the cytoskeleton post-bTBI in our study may imply a reduced ability to recover from deformation, which may be one of the factors contributing to the long-lasting behavioral and structural effects observed. In other words, this loss of elasticity to the cytoskeletal structure may impact how neurons and their glial cells maintain their morphology and connectivity in post-injury conditions and further exacerbate symptoms and impede recovery. The above studies show that the cellular and extracellular adaptations to mechanical load are with regard to the responsiveness of the brain to injury, modifying further its vulnerability to additional trauma, and pathophysiologically progression after TBI.

### 2.4 Viability and Differentiation of AHPCs After Shockwave Exposure

#### 2.4.1 Single Shockwave Exposure from ‘Top-to-Bottom’

Cell viability assessments were performed to determine if the application of the shockwave exposure influenced the cell health of the exposed cultures. The percentage of viable cells was determined using a PI staining procedure (performed 24 hours after shockwave exposure) to evaluate the short-term effect of the single shockwave exposure from top on cell health. The Propidium iodide (PI) staining performed 24 hours after shockwave exposure showed that exposure to a single shockwave blast had little to no effect on cell viability shortly after shockwave as there were little to no cells stained with PI **Fig**.A2(A-B)). The percentage of PI-labeled cells in the shockwave exposed samples was below 1% for both the control and shockwave exposed samples. The percentage of viable cells within the cultures was 99.85±0.1% for the control condition while the shockwave exposed condition had 99.68±0.1% viable cells **Fig**.A2(C). While evaluating the cell viability of the samples at the end of the culture period, the percentage of PI-labeled cells present on the coverslips remained low for both shockwave groups at 7 DIV with cell viability calculated to be 99.88±0.1% and 99.81±0.1% for the control and shockwave exposed conditions, respectively **Fig**.A3 (A-C). These results indicated that there was a negligible effect of single shockwave exposure from ‘top to bottom’ direction on the viability of the AHPCs shortly after exposure or after a longer-term culture with cell viability remaining high.

Immunocytochemistry (ICC) was used to evaluate the effect of shockwave exposure on AHPC proliferation and differentiation. The results from the immunocytochemical analysis determined that while there was a slight decrease in expression of all the antibodies tested following single shockwave exposure from the ‘top to bottom’ direction, there were no significant differences in the expression of any of the antibodies tested between the control and shockwave exposed samples (**Fig**.6(A-F’)). There was minimal change in Ki67 expression with the percentage remaining at 2.2±2.5% and 2.08±1.4% for the control and single shockwave conditions, respectively **Fig**.6(G). These results showed that a small subpopulation of the AHPCs remained proliferative by the end of the culture period, and the single shockwave exposure did not affect the proliferative ability of the AHPCs. Similarly, the percentage of TuJ1-expressing cells was 22.29±15.48% compared to 18.84±14.03% for the control and single shockwave exposure conditions, respectively, indicating little effect of the shockwave exposure on the ability of the AHPCs to differentiate into immature neurons **Fig**.6(G). MAP2ab expression showed a similar decrease in expression following single shockwave exposure from 12.54±9.15% for the control condition to 9.28±11.48% for single shockwave exposure conditions **Fig**.6(G). These results confirm the minimal effect of the single shockwave exposure from ‘top to bottom’ direction on neuronal differentiation with no significant changes in the percentage of maturing neurons in the AHPC cultures. Next, RIP expression slightly decreased, going from 39.4±16.04% for the control condition to 37.29±17.21% for the shockwave condition indicating little effect of the shockwave exposure on oligodendrocyte differentiation **Fig**.6(G). Lastly, there was no GFAP expression seen for either the control or shockwave exposed samples, indicating no astrocytes were present in the cultures with or without shockwave exposure (data not shown). Overall, these results show that single shockwave exposure from ‘top to bottom’ direction had little to no effect on cell behavior in regard to cell viability, proliferation, or differentiation.

**Fig. 6.**
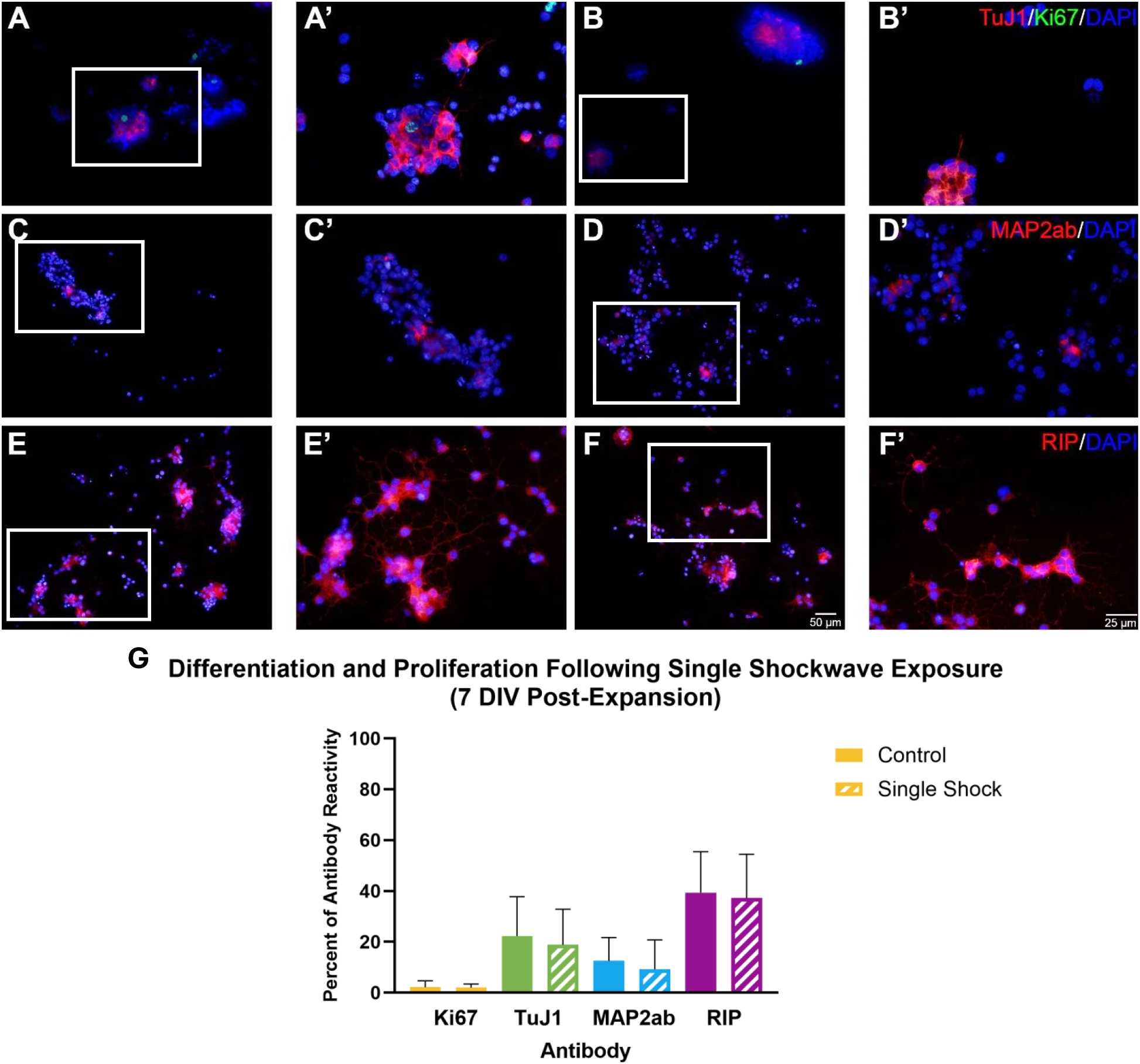
Differentiation and Proliferation of AHPCs at 7 DIV Post-Expansion Following Single Shockwave Exposure from Top to Bottom. Fluorescence images of AHPCs at 7 days *in vitro* (DIV) post-expansion following single shockwave exposure from above. Cells were single or double immunostained with an immature neuron marker (TuJ1, red; A-B’), a cell proliferation marker (Ki67, green; A-B’), a maturing neuronal marker (MAP2ab, red; C-D’), an oligodendrocyte marker (RIP, red; E-F’), and a cell nuclei marker (DAPI, blue; A-F’). Images A-F were taken using a 20x objective, with a scale bar of 50 µm. Images A’-F’ represent the boxed region in each respective image at a higher magnification, with a scale bar of 25 µm.**G:** There was no significant difference between the culture conditions. Bars represent the mean percentage of immunolabeled cells, and the error bars represent the standard deviation (± SD). *N* = 2 independent experiments, with 20 image fields quantified for each condition.

#### 2.4.2 Double Shockwave Exposure from ‘Top-to-Bottom’

PI staining was performed at both 24 hours post-exposure and at the end of the culture period to evaluate the effect of double shockwave exposure from ‘top to bottom’ direction on AHPCs. The cell viability results performed 24 hours post-exposure showed no significant change in the percentage of viable cells following double shockwave exposure **Fig**.A2(D-E). Cell viability for both groups remained above 99% with specific values of 99.89±0.04% and 99.03±0.2% for the control and shockwave exposed conditions, respectively **Fig**.A2(F). Similarly, at the end of the culture period, there was no significant difference in the percentage of viable cells between the culture conditions (99.17±0.3% compared to 99.8±0.4%) **Fig**.A3 (D-F). Taken together, these results confirmed that even after double shockwave exposure, there was no significant effect of shockwave exposure from ‘top to bottom’ direction on the cell viability of the AHPCs.

In contrast to the single shockwave exposure from top results, there were significant changes seen for both immature neuron differentiation as well as oligodendrocyte differentiation following double shockwave exposure as illustrated by the significant changes in antibody expression seen for the TuJ1 and RIP antibodies, respectively **Fig**.7(A-B’, E-F’). TuJ1 expression significantly decreased from 11.41 ± 1.7% for the control condition to 6.42 ± 0.7% for the double shockwave condition **Fig**.7(G). RIP expression showed an even greater significant decrease in expression with the average percentage of RIP-expressing cells dropping from 44.06*±*3.3% to 31*±*2.4% following double shockwave exposure from above **Fig**.7(G). Similar to the single shockwave exposure experiments, MAP2ab and Ki67 expression, while slightly decreased following double shockwave exposure, though were not significantly different from the control condition **Fig**.7(A-B’, C-D’). MAP2ab expression decreased by less than 1.5% between the control and double shock exposure conditions (11.49 ± 1.3% compared to 10 ± 1.3%) **Fig**.7(G). Ki67 expression remained around 3% for both the control and double shockwave conditions, with the specific percentage of antibody expressions being 3.44 ± 0.5% and 3.11 ± 0.4%, respectively **Fig**.7(G). Also, similar to the single shockwave exposure from the top-to-bottom experiments, no GFAP expression was detected for any of the double shockwave replicates for either the control or shockwave-exposed samples (data not shown).

**Fig. 7.**
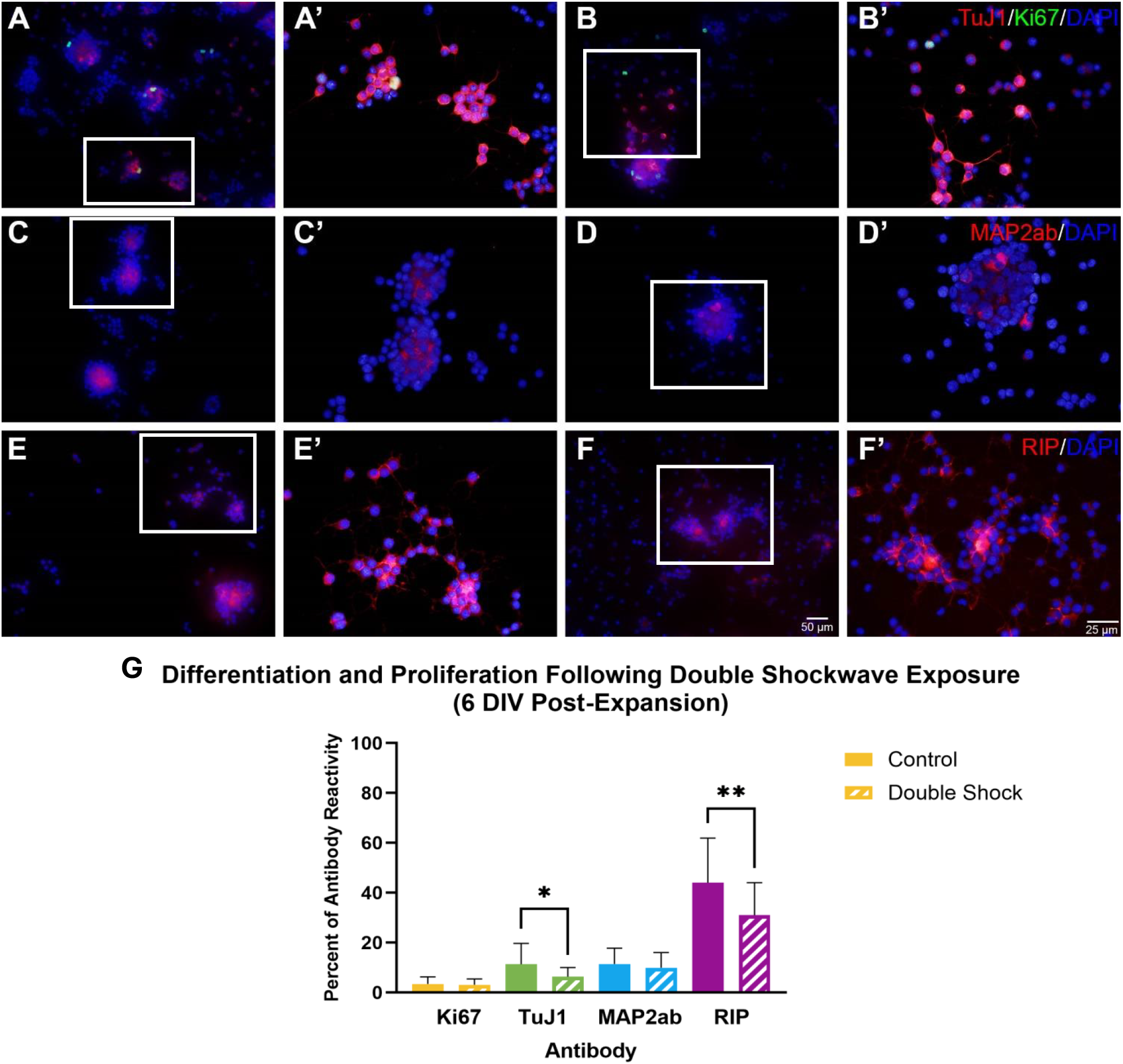
Differentiation and Proliferation of AHPCs at 6 DIV Post-Expansion Following Double Shockwave Exposure from Top to Bottom. Fluorescence images of AHPC at 6 DIV post-expansion following double shockwave exposure from ‘top to bottom’ direction. Cells were single or double immunolabeled with an immature neuron marker (TuJ1, red; A-B’), a cell proliferation marker (Ki67, green; A-B’), a maturing neuron marker (MAP2ab, red; C-D’), an oligodendrocyte marker (RIP, red; E-F’), and a cell nuclei marker (DAPI, blue; A-F’). Images A-F were taken using a 20x objective with scale bar = 50 µm. Images A’-F’ are of the boxed region in each respective image taken at higher magnification, scale bar = 25 µm. G: There was a significant decrease in the expression of TuJ1 and RIP following double shockwave exposure. There was no significant difference between the culture conditions for the remaining antibody labeling. Bars represent the mean percentage of antibody-labeled cells, and the error bars represent the standard error of the mean (±SEM). N=3 independent experiments, 30 image fields were quantified for each condition.

While the double shockwave exposure had some effect on neuronal and oligodendrocyte differentiation, overall, there was a minimal effect of the double shockwave exposure from the ‘top to bottom’ direction on cellular behavior. Cell viability remained high for both the control and shockwave exposed samples at both 24 hours post-exposure and at the end of the culture period. The double shockwave exposure from ‘top to bottom’ direction did significantly decrease the percentage of immature neurons and oligodendrocytes present within the cultures. However, double shockwave exposure had no effect on the percentage of maturing neurons or on the amount of proliferation occurring within the samples.

While comparing the results of the cell viability analyses between the single shockwave and double shockwave exposure from top-to-bottom experiments, we found that there were some significant differences in the percentage of viable cells between the conditions. At the early point, there was a significant decrease in the percentage of viable cells for the double shockwave exposed samples compared to the single shockwave exposed samples **Fig**.A4(A). However, it should be noted that the percentage of viable cells in both conditions remained above 99%. Similar results were seen for the control and shockwave exposed conditions at the later time points, with significant decreases in cell viability being found for the double shockwave conditions, though the viability of all groups remained above 98% **Fig**.A4(B). Taken together, these findings indicated that while there may have been significant differences in viability between the single and double shockwave exposed groups, the viability remained high for all conditions at both time points tested. Meanwhile, when comparing the antibody expression results of the single shockwave exposed to the double shockwave exposed samples, we found that only one antibody showed a significant change in expression between the two exposure conditions. The expression of TuJ1, the immature neuron marker, was significantly lower in the double shockwave exposed samples compared to the single shockwave exposed samples (18.84±14.03% compared to 6.42±0.7%) **Fig**.A4(C). No other antibodies showed any significant change in expression between the shockwave exposure conditions. In summary, single shockwave exposure from the ‘top to bottom’ direction did not have an effect on cell behavior in terms of cell viability, proliferation, or differentiation. However, for double shockwave exposures, significant changes were observed between the exposed and control conditions. Specifically, significant changes in the expression of the immature neuron and oligodendrocyte markers became evident. When we compared the differences in cell viability and differentiation between the shockwave conditions, a significant decrease in cell viability was determined following double shockwave exposure; however, the cell viability remained high for all conditions. Lastly, a significant decrease in immature neuron expression was observed following double shockwave exposure in comparison to a single shockwave exposure.

#### 2.4.3 Single Shockwave Exposure from ‘Bottom-to-Top’

Cell viability following below shockwave exposure was assessed 3 hours post-shockwave exposure to evaluate the short-term effects of a single shockwave exposure on cell viability. Results indicated that exposure to a single shockwave blast administered below the dish had minimal to no effect on cell viability, as little to no remaining adherent cells were stained with PI **Fig**.A5(A-B). The percentage of PI-labeled cells was less than 1% for the control and below-shockwave exposed samples, with viable cell percentages of 99.45±0.2% for the control and 99.6±0.2% for the treated samples **Fig**.A5(C). Similarly, at the end of the extended culture period (4 DIV), less than 1% of cells on the coverslips were positively labeled with PI. The percentage of viable cells remained at 99.21±0.3% for the control condition and 99.45±0.2% for the shockwave-exposed cells **Fig**.A6(A-C). The cell viability results at both the 3-hour post-shockwave exposure and 4 DIV postexpansion time points showed no significant differences between shockwave-exposed samples and non-exposed controls. The percentage of PI-labeled cells at both time points remained below 1%, confirming that the single shockwave blast from below did not impact cell viability.

As observed in the previous treatments, the percentage of AHPCs at 4 DIV post-expansion following a single shockwave exposure from ‘bottom to top’ direction showed no significant changes in antibody expression between the control and shockwave-exposed samples. There was also a lack of a consistent trend regarding the effects of the below shockwave exposure on AHPC differentiation and proliferation **Fig**.8(A-F’). Specifically, both TuJ1 and Ki67 showed slightly decreased expression in the shockwave-exposed samples compared to the control, with a decrease from 20.05 ± 1.6% to 18.83 ± 1.8% and 3.38 ± 0.7% to 3.25 ± 0.6%, from the control to the shockwave condition, respectively**Fig**.8 (G). This indicated that experiencing a single shockwave from ‘bottom to top’ direction had minimal effect on the ability of AHPCs to differentiate into immature neurons or their ability to proliferate. Conversely, MAP2ab antibody expression increased from the control at 12.85 ± 1.1% to 13.79 ± 0.8% for the shockwave-exposed cells, confirming the lack of effect of the single shockwave from ‘bottom to top’ direction on neuronal differentiation, with no significant changes in the percentage of maturing neurons within the culture **Fig**.8(G). The positive expression of the RIP antibody followed a similar trend, with a slight increase from 21.21 ± 1.19% to 24.17 ± 1.8% between the control and shockwave-exposed cultures **Fig**.8(G). This demonstrates that the shockwave had little effect on the ability of the AHPCs to differentiate into mature neurons. No GFAP-immunoreactive AHPCs were detected, indicating no astrocyte differentiation was present in the cultures (data not shown). Although TuJ1 and Ki67 expression exhibited slight decreases post-exposure, and MAP2ab and RIP showed minimal increases, the lack of a consistent trend regarding the effects of a single shockwave (administered from ‘bottom to top’ direction) on the AHPCs reinforces the notion that there were no significant changes in differentiation or proliferation of the cells.

**Fig. 8.**
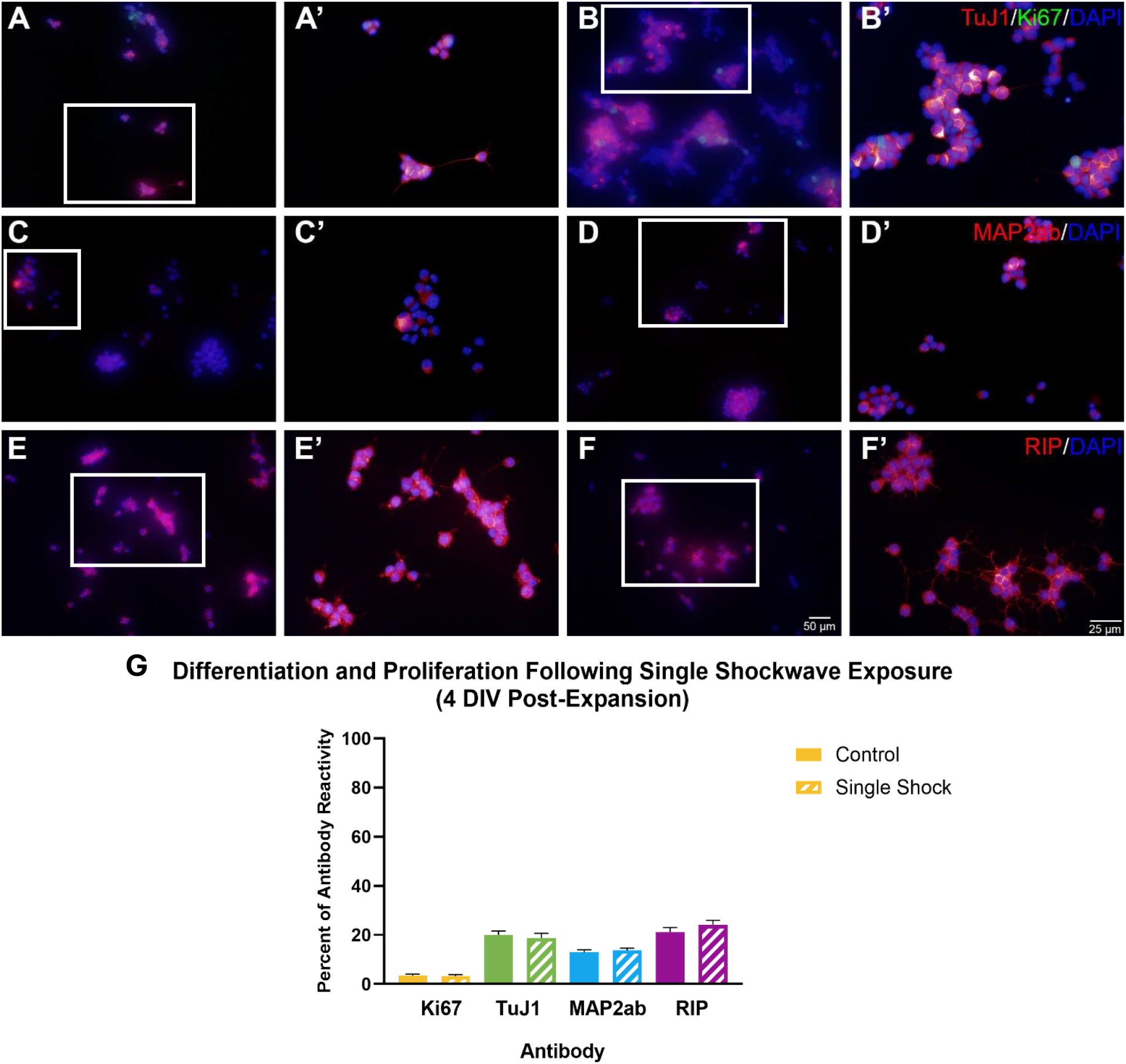
Differentiation and Proliferation of AHPCs at 4 DIV Post-Expansion Following Single Shockwave Exposure from Bottom to Top. Fluorescence images of AHPCs at 4 DIV post-expansion following single shockwave exposure from below. Cells were single or double immunostained with an immature neuron marker (TuJ1, red; A-B’), a cell proliferation marker (Ki67, green; A-B’), a maturing neuron marker (MAP2ab, red; C-D’), an oligodendrocyte marker (RIP, red; E-F’), and a cell nuclei marker (DAPI, blue; A-F’). Images A-F were taken using a 20x objective with scale bar = 50 µm. Images A’-F’ are of the boxed region in each respective image were taken at higher magnification, scale bar = 25 µm. G: There was no significant difference between the culture conditions. Bars represent the mean percentage of antibody-labeled cells, and the error bars represent the standard error of the mean (±SEM). N=3 independent experiments, 30 image fields were quantified for each condition.

#### 2.4.4 Double Shockwave Exposure from ‘Bottom-to-Top’

Analysis of cell viability 3 hours after shockwave exposure indicated that a double shockwave application from ‘bottom to top’ direction had minimal impact on the ability of AHPCs to survive in culture, as there were no significant changes in the amount of PI staining between the conditions **Fig**.A5(D-E). Quantitative analysis showed that the percentage of PI was below 1% for both the control and shockwave-treated samples, with 98.55±0.5% and 98.73±0.5% viable cells, respectively **Fig**.A5(F). Similarly, evaluation of cell viability at 4 DIV post-expansion showed less than 1% cell death, with 99.26±0.3% viable cells in the control condition and 99.812±0.3% in the shock condition **Fig**.A6(D-F). Cumulatively, these results indicated that double shockwave exposure from ‘bottom to top’ direction had little to no effect on the viability of remaining adherent AHPCs, both shortly after the shockwave exposure or following expansion and long-term culture, as cell viability remained consistently high in both conditions.

Quantification of the immunocytochemical analysis revealed a consistent slight increase in the expression of all antibodies tested following double shockwave exposure from ‘bottom to top’ direction; however, no significant differences were observed between the control and shockwave-exposed samples **Fig**.9(A-F’). There was a slight change in Ki67 expression between the control and shockwave-treated samples, increasing from 2.69±0.3% to 3.1±0.3% of positively labeled cells **Fig**.9(G). This suggests that a subpopulation of AHPCs remained proliferative in both conditions, indicating that the shockwave did not affect their ability to divide. Similarly, TuJ1-immunoreactivity, showed an increase from 29.12±1.8% in the control group to 32.01±2.6% in the shockwave-exposed cultures **Fig**.9(G). This indicated that the shockwave had a limited impact on the cells’ ability to differentiate into immature neurons. MAP2ab expression also followed this trend, increasing slightly from 15.4±1.2% to 15.61±1.2% from the control to the shockwave condition, respectively **Fig**.9(G). These results confirmed that double shockwave exposure from below had minimal impact on neuronal differentiation, with no significant changes in the percentage of immature or maturing neurons within the AHPC cultures. The expression of RIP increased from 20.18±1.3% to 22.52±1.2% between the control and shockwave-treated samples, indicating the limited effect of double shockwave exposure from ‘bottom to top’ direction on oligodendrocyte differentiation **Fig**.9(G). Consistent with the results for the single shockwave from the ‘bottom to top’ direction results, no GFAP expression was observed within any culture condition (data not shown). Overall, while there was a consistent pattern of slight increases in antibody expression following double shockwave exposure from ‘bottom to top’ direction, these results demonstrated no significant differences between the control and shockwave-exposed samples, further indicating a lack of impact on cell behavior in terms of viability, proliferation, or differentiation.

**Fig. 9.**
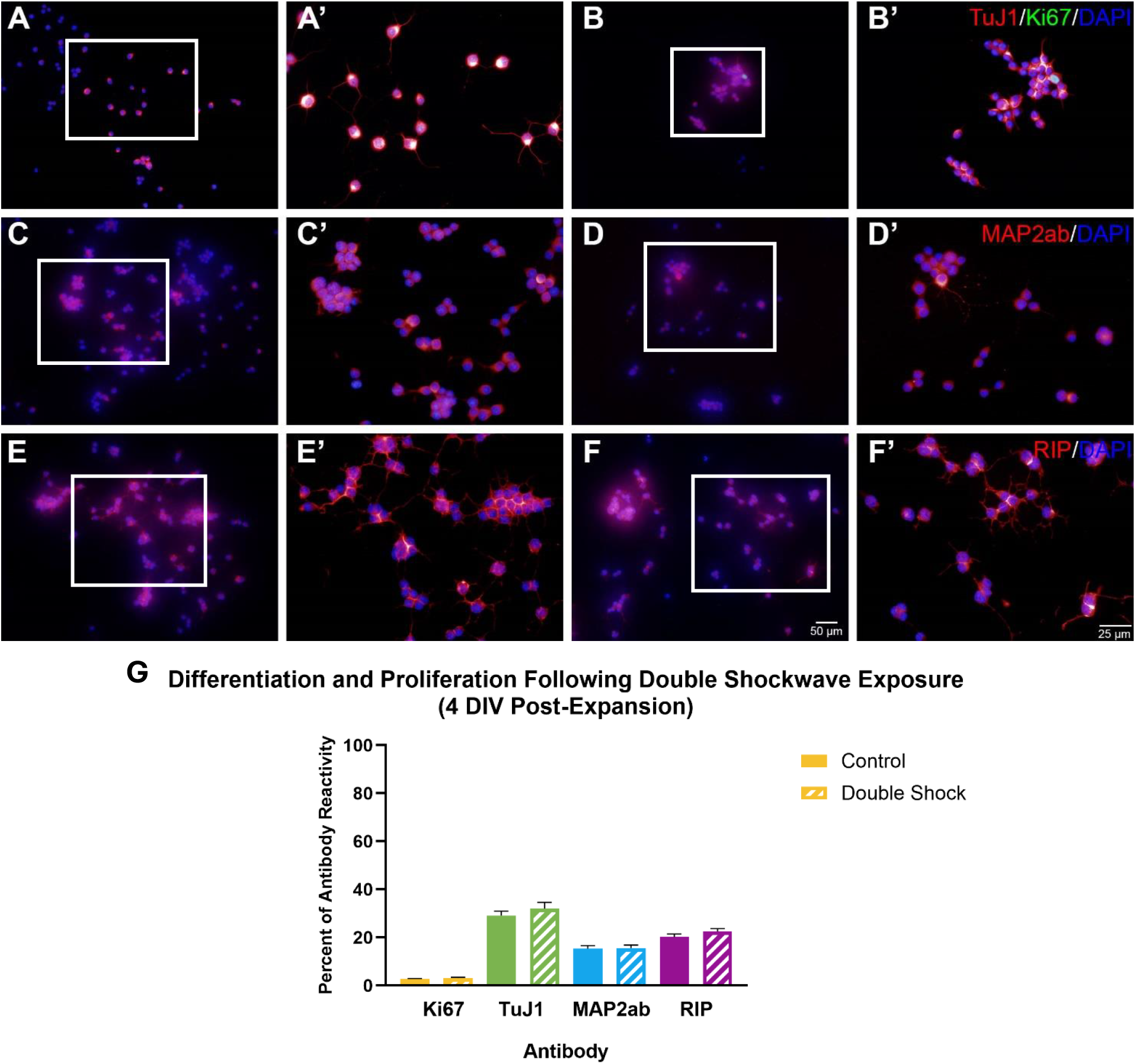
Differentiation and Proliferation of AHPCs at 4 DIV Post-Expansion Following Double Shockwave Exposure from Bottom to Top. Fluorescence images of AHPCs at 4 DIV post-expansion following single shockwave exposure from below. Cells were single or double immunostained with an immature neuron marker (TuJ1, red; A-B’), a cell proliferation marker (Ki67, green; A-B’), a maturing neuron marker (MAP2ab, red; C-D’), an oligodendrocyte marker (RIP, red; E-F’), and a cell nuclei marker (DAPI, blue; A-F’). Images A-F were taken using a 20x objective with scale bar = 50 µm. Images A’-F’ are higher magnification images of the boxed region in each respective image, scale bar = 25 µm. G: There was no significant difference between the culture conditions. Bars represent the mean percentage of antibody-labeled cells, and the error bars represent the standard error of the mean (±SEM). N=3 independent experiments, 30 image fields were quantified for each condition.

When comparing the cell viability results from the single shockwave and double shockwave below exposure experiments, it was found that there were no significant differences in the percentage of PI-labeled cells **Fig**.A7(A). When evaluating the results from the early time point, we saw a slight increase in the amount of PI-labeling following double shockwave exposure conditions in both the control and exposed conditions compared to a single shockwave exposure. Cell viability was 99.45±0.2% for the single shockwave samples and 98.55±0.5% for the double shockwave samples for the control condition **Fig**.A7(A). Meanwhile, the single shockwave condition had a viability of 99.6±0.2% that decreased to 98.73±0.5% following double shockwave for the exposed samples **Fig**.A7 (A). Furthermore, the same pattern was seen following the PI staining performed at the end of the culture period. Here, the single shockwave control had a cell viability of 99.21±0.3%, whereas the double shockwave control viability decreased slightly to 99.26±0.3% **Fig**.A7(B). The single shockwave exposed sample had a viability of 99.45±0.2%, that again decreased to 99.812±0.3% in the double shockwave exposed sample **Fig**.A7(B). Taking these results together, not only were the control and shockwave exposure conditions not significantly different from each other, but the single and double shockwave conditions showed no significant differences between each other for either culture condition. These findings indicated that increasing the amount of shockwave exposures did not affect cell viability when the shockwave was applied from ‘bottom to top’ direction on the dishes. Recall that when the shockwave exposure was applied from above’top to bottom’ direction, it was found that there was a significant decrease in cell viability following double shockwave exposure compared to the single shockwave exposure application. However, it is important to note that the average percentage of PI-labeled cells for all conditions, single and double shockwave exposure for both control and shockwave exposed groups, were below 2%. When comparing the ICC results between the single and double shockwave exposure from below experiments, it was found that double shockwave exposure resulted in significantly higher TuJ1 expression, with 32.01±2.6% of cells showing positive immunolabeling, compared to 18.83±1.8% for single shockwave exposure condition **Fig**.A7(C). However, no significant differences or patterns were observed in the expression of the other antibodies tested. For instance, MAP2ab expression increased from 13.79±0.8% after single shock exposure to 15.61±1.2% after double shock exposure **Fig**.A7(C). In contrast, RIP showed a nonsignificant decrease, with positive expression dropping from 24.17±1.8% after single shock to 22.52±1.2% after double shock **Fig**.A7(C). Ki67 exhibited a similar trend, decreasing from 3.25±0.6% positive labeling after single shock to 3.1±0.3% after double shock **Fig**.A7(C). These results suggest that a minimum of double shock is required to observe a significant effect on AHPC cell behavior.

In summary, for all shockwave conditions performed, no significant changes in cell viability occurred at either the early or later time points. For all conditions, the viability remained high (98.5%). These findings are consistent with findings from other groups applying a similar or high-strength shockwave blast using animal or organoid models [56, 57]. It should be noted that the viability analyses performed in the current study were unable to account for any cells that may have been lost due to detachment as a result of the shockwave blast exposure. Meaning, the cell viability values obtained for this study are the viability percentages for the cells that remained adhered to the substrates post-shockwave exposure. Similarly, we found that the shockwave blast exposure had little effect on AHPC cell proliferation or differentiation, which is consistent with the findings of other groups who used similar or slightly greater shockwave blast conditions [57, 58]. Only following double shockwave exposure from top were significant changes in differentiation observed. Specifically, there was a significant decrease in the percentage of TuJ1 and RIP-expressing cells, indicating a decrease in the amount of immature neurons and oligodendrocytes in the cultures following a higher magnitude of shockwave blast exposures. Previous findings from our group showed the percentage of maturing neurons and oligodendrocytes present in AHPC neurosphere cultures decreased in a near stepwise fashion with increasing shockwave strength [59]. The current results appear to agree with these findings in that while the shockwave strength remained constant, increasing the number of blasts may have a similar negative effect on AHPC cell differentiation. Interestingly, when the shockwave blast was oriented below the culture dish for ‘bottom to top’ double shockwave exposures, there was a slight increase in the percentage of TuJ1 expressing cells. It was also found that the double shockwave exposure from below (from ‘bottom to top’ direction) had a significantly h igher a mount o f T uJ1 e xpression c ompared t o a s ingle shockwave exposure. One explanation for the increase in the amount TuJ1 present in these cultures could be that some level of neurogenesis is occurring following shockwave exposure in an effort to offset the loss of neurons due to increased shockwave injury that occurred when the blast was coming from below the dish as opposed to above the dish [60].

Lastly, it is important to note the potential discrepancies in the significant changes following shockwave exposure between the results collected via AFM analysis and the immunocytochemical analyses. The results collected from the AFM analysis were collected within hours of the shockwave blast exposure with samples being analyzed minutes after the application of the blast. The immediate analyses of these samples would not allow cells injured by the blast exposure to recover, thus resulting in more significant differences between the exposed samples and the non-exposed control. In contrast, the immunocytochemical analyses were not performed until days later, after the cells had a chance to recover from the blast exposure. The additional growth period the cells experienced following shockwave exposure likely allowed cells that might have been injured post-exposure to sufficiently re cover to a si milar st ate as th e no n-exposed co ntrol sa mples by the time immunocytochemical analyses occurred resulting in the non-significant d ifferences in di fferentiation or proliferation between these groups at the time of analysis.

### 2.5 Impact of Shockwave Propagation Direction

Impedance matching controls how much of the shockwave is reflected or transmitted at each interface during shockwave propagation across a multitude of different materials. Since wave impedance is simply the product of the material’s density and wave speed, a transition at an interface from high to low or from low to high impedance will affect the energy distribution in the wave and thus determine both reflected and transmitted wave intensities. That difference, in fact, can completely change the effectiveness of the interaction between the shockwave and cellular structures. For example, in a top-to-bottom setup, each layer has its appropriate impedance mismatch, reflecting waves that weaken the shockwave and thus result in minimal disruption visible in cytoskeletal structures. As indicated by Zhao et al. [61], the reflection and transmission of the shock waves at interfaces (described by the multi-impedance materials) depend upon the following equations from, given as:

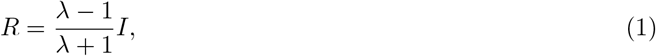

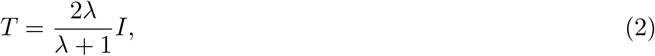

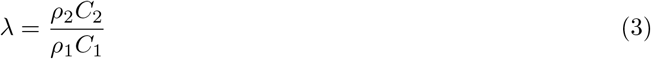

where *λ* denotes the wave impedance ratio, with *ρ*_1_ and *ρ*_2_ as the densities of materials 1 and 2, respectively, and *C*_1_ and *C*_2_ as the corresponding wave speeds in each material.

This could suggest that top-to-bottom shockwaves cannot be deep enough to penetrate and cause significant disturbance to the intracellular network because every layer of the media dissipates energy from the shockwaves. However, in the process of bottom-to-top exposure, the shockwave directly acts on the bottom of the petridish with only one impedance mismatch; thus, a stronger and more concentrated force could be delivered to the cells, with much lesser dissipation of energy. The configuration that was seemingly designed to enhance shear forces on the cytoskeleton through more pronounced morphological changes would indicate more disturbance within the cellular nanostructure. The impedance matching [61] and energy dissipation principle [62] apparently support these views, as transitions of impedance are fewer, thus allowing for an easy transfer of shockwave energy into the cells, leading to observable shifts within the cellular shape and structural integrity.

## 3 Conclusion

In the present study, shockwaves of different magnitudes, durations, and directions were applied to AHPCs to study the structural, nanomechanical, and viscoelastic properties of cells as well as cell survival, differentiation, proliferation using AFM and ICC assays, respectively. The results from AFM suggested that the bottom-to-top shockwave exposures had a greater disruptive effect on the actin cytoskeletal network, inducing severe changes in cell morphology than the top-to-bottom direction of exposure. With increased intensity of shockwaves and direct impact through one polystyrene layer of the cell petridish, the cells became more rounded within one hour of exposure. Nanomechanical property analyses showed that cells became softer and more deformable with the increased magnitude of shockwaves. This softening and deformation were even more dramatic in the case of direct bottom-to-top shockwaves. The cells also showed lower viscosity and shorter relaxation times, managing to regain their original shape faster after being exposed to a higher magnitude of shockwave. Although such structural changes were observed, PI staining results showed that cell viability remained consistently high throughout all conditions of shockwave exposures. However, it has to be noted that these viability values only correspond to the surviving cells that remained adhered to the substrates post-exposure, since the detached cells were not accounted for. Shockwave exposure also demonstrated limited effects on AHPC cell proliferation and differentiation under most conditions; however, there was a significant decline in TuJ1 and RIP expression after double shockwave exposure from ‘top-to-bottom’ direction, which indicates a reduced population of immature neurons and oligodendrocytes respectively. Conversely, double shockwave exposure from ‘bottom-to-top’ direction resulted in a slight increase in TuJ1 expression that may indicate neurogenesis as a result of compensatory response after injury from higher intensity blast, crossing the overpressure magnitude threshold inside cells. Notably, it was realized that there were some differences between AFM and ICC results that might be attributed to the timing when the two kinds of analyses were performed. The AFM measurements taken shortly after exposure to shockwaves showed immediate responses in the cells, such as highly significant structural changes, while ICC analyses involved putting the shockwave-exposed cells back in the incubator, providing the surviving cells a chance to recover. These findings underline that the responses of cells to mechanical trauma are dynamic and time-dependent and illustrate the importance of integrating multiple approaches of analysis for capturing immediate and long-term effects of shockwave exposure. This study is the first of its kind to explore the correlation among cellular differentiation and cellular nanomechanics post bTBI.

## 4 Materials and Methods

### 4.1 Cell Culture

Adult rat hippocampal progenitor cells (AHPCs) were gifted by Dr. F.H. Gage, Salk Institute, La Jolla, CA. AHPCs were cultured in a T25 tissue culture flask (Thermo Fisher Scientific) coated with poly-L-ornithine (10 µg/mL; Sigma Aldrich) and laminin (10 µg/mL; Cultrex by Trevigen). Cells were cultured in maintenance medium (MM) composed of Dulbecco’s modified Eagle’s medium/Ham’s F-12 (DMEM/F-12, 1:1; Gibco by Thermo Fisher Scientific) supplemented with 2.5 mM GlutaMAX (Thermo Fisher Scientific), 1 x N2 supplement (Gibco by Thermo Fisher Scientific),1 x Penicillin/streptomycin (Gibco by Thermo Fisher Scientific), and 20 ng/mL basic fibroblast growth factor (bFGF; Gibco by Thermo Fisher Scientific). The cells were incubated at 37 °C in a 5% CO_2_ atmosphere. AHPCs in the flask were fed with MM every 2 days by performing half media changes. They were harvested when the cell confluency reached 80%.

#### Top-to-Bottom Shockwave Exposure Experiments

Cells were initially seeded into poly-L-ornithine/laminin (POL) coated 35 mm petridishes (Falcon) at a density of 40,000 cells/dish. Dishes were maintained in MM for 6 days *in vitro* (DIV) with standard half media changes. At 4 DIV, a subset of the dishes was exposed to shockwave blast injury. The cells within the dishes were cultured for an additional 48 hours at which time the cells from the dishes were harvested and expanded onto POL-coated coverslips at a density of 3,000 cells/coverslip. Half of the coverslips were seeded with non-exposed cells while the remaining half were seeded with cells exposed to the shockwave blast. Coverslips were cultured in MM for the first 24 hours after which time the media was switched to differentiation medium (DM; MM without the bFGF), and the cells were maintained in DM until the end of the culture period. For the single shockwave blast exposure experiments the coverslips were maintained until 13 DIV, and for the double shockwave blast exposure the coverslips were maintained until 12 DIV (see Supplementary **Fig**.A1).

#### Bottom-to-Top Shockwave Exposure Experiments

Cells were initially seeded into POL-coated 35 mm petridishes at a density of 20,000 cells/dish. Dishes were maintained in MM for 6 DIV with standard half media changes. At 4 DIV, a subset of the dishes was exposed to shockwave blast injury. The cells within the dishes were cultured for an additional 48 hours at which time the cells from the dishes were harvested and expanded onto POL-coated coverslips at a density of 2,000 cells/coverslip. Half of the coverslips were seeded with non-exposed cells while the remaining half were seeded with cells exposed to the shockwave blast. Coverslips were cultured in MM for the first 2 4 h ours a nd t he m edia w as s witched t o D M a fter t hat. The cells were maintained in DM until the end of the culture period. For both the single and double shockwave exposure experiments the coverslips were maintained until 10 DIV (see Supplementary **Fig**.A1).

### 4.2 Atomic Force Microscope (AFM) Setup

Nanomechanical measurements of the AHPCs were performed using an atomic force microscope (AFM) (Bruker BioScope Resolve system) integrated with a NanoScope V controller. The AFM was operated in Peak Force Quantitative Nanomechanical Mapping (PF-QNM) mode, which simultaneously provided high-resolution imaging and nanomechanical property mapping. This mode enabled precise measurements of key mechanical parameters such as Young’s modulus, deformation, dissipation, and adhesion, all while minimizing the risk of damaging the delicate biological samples. Immediately after shockwave exposure, the dish with AHPCs was placed on the AFM stage (connected to the sample heater) in a vibration isolation chamber to maintain a controlled temperature environment during measurements. Pre-calibrated PFQNM-LC-CAL-A probes, specifically designed for live cell applications, were employed for single-cell measurements. These probes, with a 70 nm tip radius and a nominal spring constant ranging from 0.074 N/m to 0.101 N/m, offered the necessary sensitivity and flexibility for probing soft biological materials without inducing excessive deformation. A controlled force of 300 pN was applied during each indentation, ensuring gentle yet consistent contact with the cell membrane and minimizing structural disruption. The AFM scan rate was set at 0.242 Hz, with scan sizes ranging from 17 *µ*m to 50 *µ*m, achieving a balance between spatial resolution and throughput. This allowed for accurate nanomechanical mapping of each cell’s surface topography, capturing critical mechanical properties. For viscoelastic property measurements, stress relaxation curves were generated using an MLCT-D probe in contact mode. These probes featured a tip radius of 20 nm and a spring constant of 0.03 N/m, making them suitable for capturing the time-dependent mechanical behavior of the cells. All measurements were conducted under carefully controlled ambient conditions, with temperature and humidity regulated to preserve cell viability throughout the experimental process.

### 4.3 Analysis for Nanomechanical Measurements

The nanomechanical data obtained from individual cells were processed and analyzed using Bruker NanoScope Analysis software. This software enabled detailed examination of both the structural and mechanical properties of the cells at nanoscale. High-resolution images and nanomechanical maps were exported as .spm files, providing comprehensive visualization of cell surface topography and mechanical characteristics. Force curves were captured as .pfc files at 10 random locations on each cell and analyzed using the Hertz model [63, 64] with a spherical indenter, yielding average Young’s modulus values for 20 cells to assess their stiffness and elastic properties. In the Hertz model, the indentation force is written as follows:

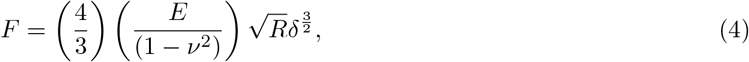

Indentation force *F*, Young’s modulus *E*, Poisson ratio *?*, indentation *δ*, and *R* representing the radius of the indenter are the parameters included in the Hertz model.

To characterize viscoelastic behavior, time-dependent force curves were obtained by maintaining a fixed position of the height sensor for two seconds during each measurement. The resulting stress relaxation curves were analyzed using Kohlrausch-Williams-Watts (KWW) models [65] to extract parameters such as retardation time and viscosity described by the following expression:

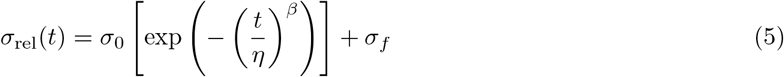

The final stress is represented by *σ*_*f*_ as time (*t*) goes towards infinity. Likewise, *σ*_0_ is the function representing stress which is dependent on *t*. The shape parameter (*β*) and characteristic life (*η*) have relationships with the time that the load has been applied to. These metrics characterize the viscoelastic behavior of the material and give insight into how the stress develops when the load has been applied for a very long period of time.

A total of 40 data sets per cell type were analyzed, offering a thorough assessment of the viscoelastic properties of the cells. Representative stress relaxation curves for each condition are shown in **Fig**.5, illustrating the distinct mechanical responses of the different cell lines.

### 4.4 Propidium iodide (PI) staining

Cell viability was determined using propidium iodide (PI; Sigma Aldrich) to stain dead cells within the cultures. For the ‘top to bottom’ shockwave exposure experiments, PI staining was performed at 5 days *in vitro* (DIV) (24 hours after shockwave blast injury) and at the end of the culture period (see supplementary **Fig**.A1(A-B). For the ‘bottom to top’ shockwave exposure experiments, PI staining was performed within 6 hours following shockwave exposure and at the end of the culture period (see supplementary **Fig**.A1(C). The PI was diluted to 1.5 µM in cell culture medium. The cells were incubated in PI solution for 20 minutes at 37°C in 5% CO_2_. As a reagent control, one coverslip with cells was incubated with 70% ethanol for 5 minutes to intentionally kill all the cells prior to the addition of the PI solution. After the 20-minute incubation, the cells were rinsed with ice-cold 0.1 M PO_4_ buffer for one minute and then were fixed with 4% paraformaldehyde (PFA, Thermo Fisher Scientific) made in 0.1 M PO_4_ for 20 minutes. Cells were then rinsed three times for 7 minutes each with phosphate-buffered saline (PBS) to remove all the fixative and then incubated with 4’,6-diamidino-2-phenylindole (DAPI; 1:500; Invitrogen by Thermo Fisher Scientific) diluted in PBS for 60 minutes at room temperature in the dark. The cells were then rinsed four times for 8 minutes each with PBS. Coverslips were then mounted onto microscope slides with Fluoromount-G with DAPI (Invitrogen by Thermo Fisher Scientific).

### 4.5 Immunocytochemistry (ICC)

At the end of the culture period, immunocytochemistry was used to evaluate the extent of proliferation and differentiation of the AHPCs following shockwave exposure **Fig**.A1. Briefly, the AHPCs on the coverslips were rinsed with ice-cold 0.1 M PO_4_ buffer for one minute and then incubated in 4% paraformaldehyde (PFA) made in 0.1 M PO_4_ buffer for 20 minutes. Following fixation, the coverslips were rinsed three times for 7 minutes each with phosphate-buffered saline (PBS) to remove all the fixatives. The coverslips were then incubated in a blocking solution composed of 0.2% Triton X-100 (Thermo Fisher Scientific), 5% normal donkey serum (NDS) (Jackson ImmunoResearch), and 0.4% bovine serum albumin (BSA) (Sigma-Aldrich) in PBS at room temperature for 90 minutes. Primary antibodies—Rabbit *α* Ki-67 (1:300, IgG; Abcam), Mouse *α* TuJ1 (1:200, IgG; R&D Systems), Mouse *α* MAP2ab (1:200, IgG; Sigma-Aldrich), Mouse *α* RIP (1:200, IgG; DSHB, Iowa City, IA), and Mouse *α* GFAP (1:200, IgG; Sigma-Aldrich)—were diluted in the blocking solution. The cells were incubated in the primary antibody solution overnight at 4°C. The next day, the samples were rinsed four times for 8 minutes each with PBS. To prepare the secondary antibody solution, Donkey *α* Rabbit AF488 (1:300, IgG; Jackson ImmunoResearch), Donkey *α* Mouse AF488 (1:200, IgG; Jackson ImmunoResearch), and Donkey *α* Mouse Cy3 (1:200, IgG; Jackson ImmunoResearch) were diluted using the blocking solution, which also contained DAPI (1:500). The cells were incubated with the secondary antibody solution at room temperature in the dark for 90 minutes and then were rinsed with PBS four times for 8 minutes each. Coverslips were then mounted onto microscope slides with Fluoromount-G with DAPI.

### 4.6 Leica Fluorescent Microscope

For cells cultured in 35 mm dishes, the cells were imaged using an inverted Leica fluorescent microscope (Leica DMI4000B; Leica Microsystems) equipped with standard epifluorescence illumination and a Leica DFC310 FX (Leica Microsystems) digital camera. For cells cultured on coverslips, the cells were imaged using an upright Leica fluorescent microscope (Leica DM5000B; Leica Microsystems) equipped with standard epifluorescence illumination and a Q Imaging Retiga 2000R (Q Imaging) digital camera. A 20 × objective was used to obtain images for quantitative data analysis, while a 40 × objective was used to obtain high magnification images of cellular morphology.

#### Data acquisition and statistical analysis

The images of the AHPCs were analyzed and quantified using ImageJ software (http://imagej.nih.gov/ij). PI, Ki-67, TuJ1, MAP2ab, and RIP immunoreactive cells were counted using the Cell Counter tool in ImageJ, and the percentage of immunoreactive cells was determined as the number of positively labeled cells divided by the number of DAPI-labeled nuclei. Ten image fields were quantified for each stain or antibody for each condition. All means are reported with standard deviation (mean ± SD) or standard error of the mean (mean ± SEM). Graph Pad Prism 10 (Graph Pad Software, Inc., San Diego, CA) was used for statistical analysis and graph-making. Means were compared using an unpaired T-test with Welch’s correction or ordinary one-way ANOVA, with statistical significance determined using Tukey’s multiple comparison test, with *α* = 0.05. All AFM data were also presented as mean values with the standard deviation (mean ± SD). Statistical analyses and graph generation were performed using OriginPro 2022 (64-bit) SR1 version 9.9.0.225 (Academic). For comparisons of mean values between groups, paired sample T-tests were conducted. Statistical significance was determined using a two-tailed t-test, with a p-value of less than 0.05 considered significant. The significance levels are denoted as follows: * for *P <* 0.05; ** for *P <* 0.01; *** for *P <* 0.001; and **** for *P <* 0.0001.

## Acknowledgements

We would like to thank Rachel Currant for her contributions to the experiments, including assistance with cell culturing, staining procedures (PI and ICC), imaging, and data analysis.

## Declarations

## Ethics approval and consent to participate

Not applicable

## Data and Materials Sharing

All materials and data are included in the supplementary information and will be available publicly.

## Author contribution

Conceptualization, A.S; Methodology, N.M, C.F, B.M, M.H.H.H, W.J.J, D.C.R, C.R; Formal Analysis, N.M, C.F, B.M, M.H.H.H; Investigation, A.S, D.S.S; Writing - Original Draft, N.M, C.F, B.M, M.H.H.H; Writing - Review & Editing, N.M, C.F, B.M, M.H.H.H, W.J.J, D.C.R, C.R, S.A.B, D.S.S, A.S; Supervision, A.S, D.S.S, S.A.B

## Appendix A Supplementary Information

**Fig. A1.**
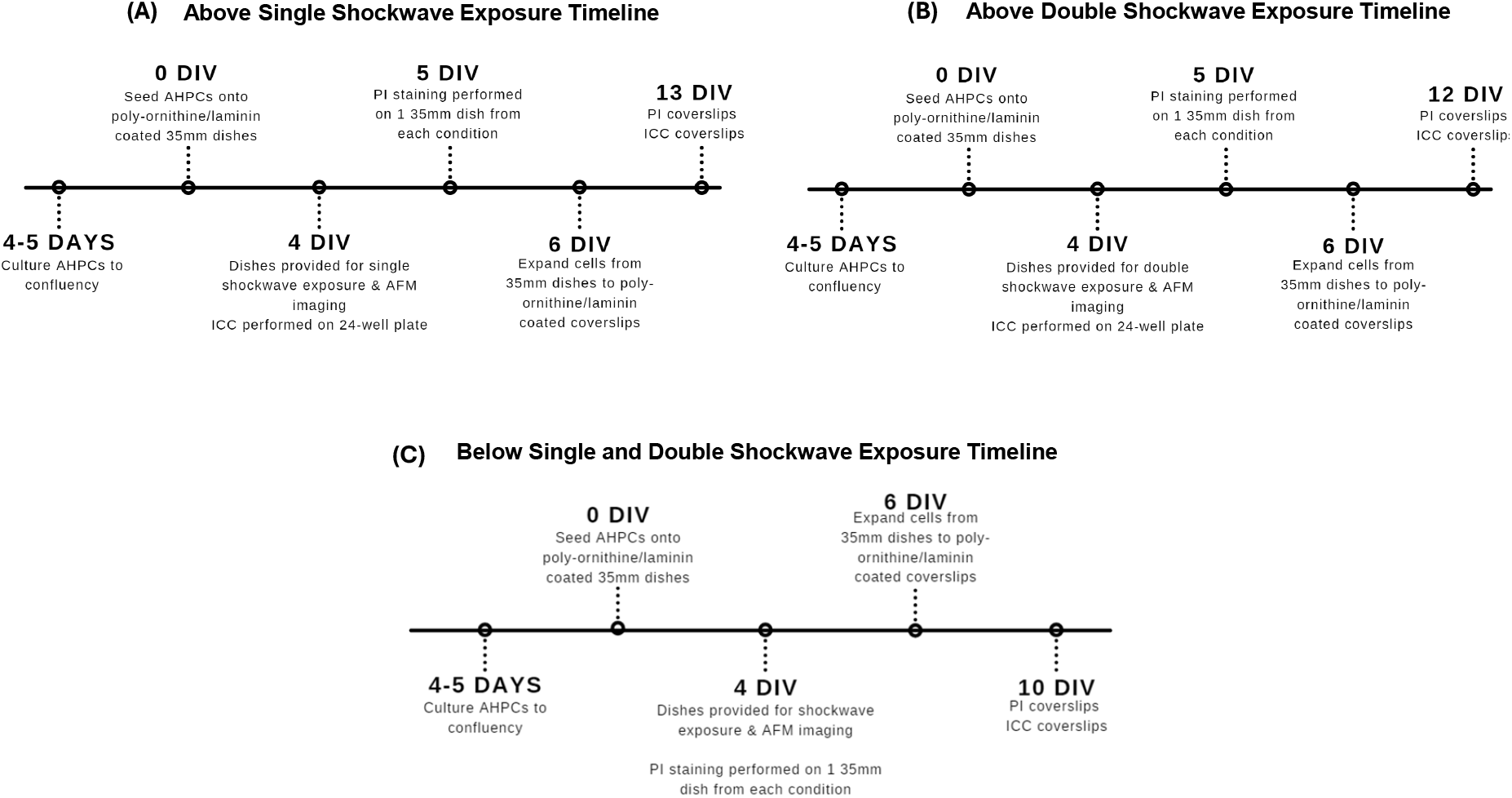
Experimental Timelines for the Top-to-Bottom and Bottom-to-Top Single and Double Shockwave Exposure Experiments. A: illustrates the culture and experimental timeline for analyses for the single above shockwave exposure samples. B: illustrates the culture and experimental timeline for analyses for the double above shockwave blast exposure samples. C: illustrates the culture and experimental timeline for analyses for the single and double below shockwave blast exposure samples.

**Fig. A2.**
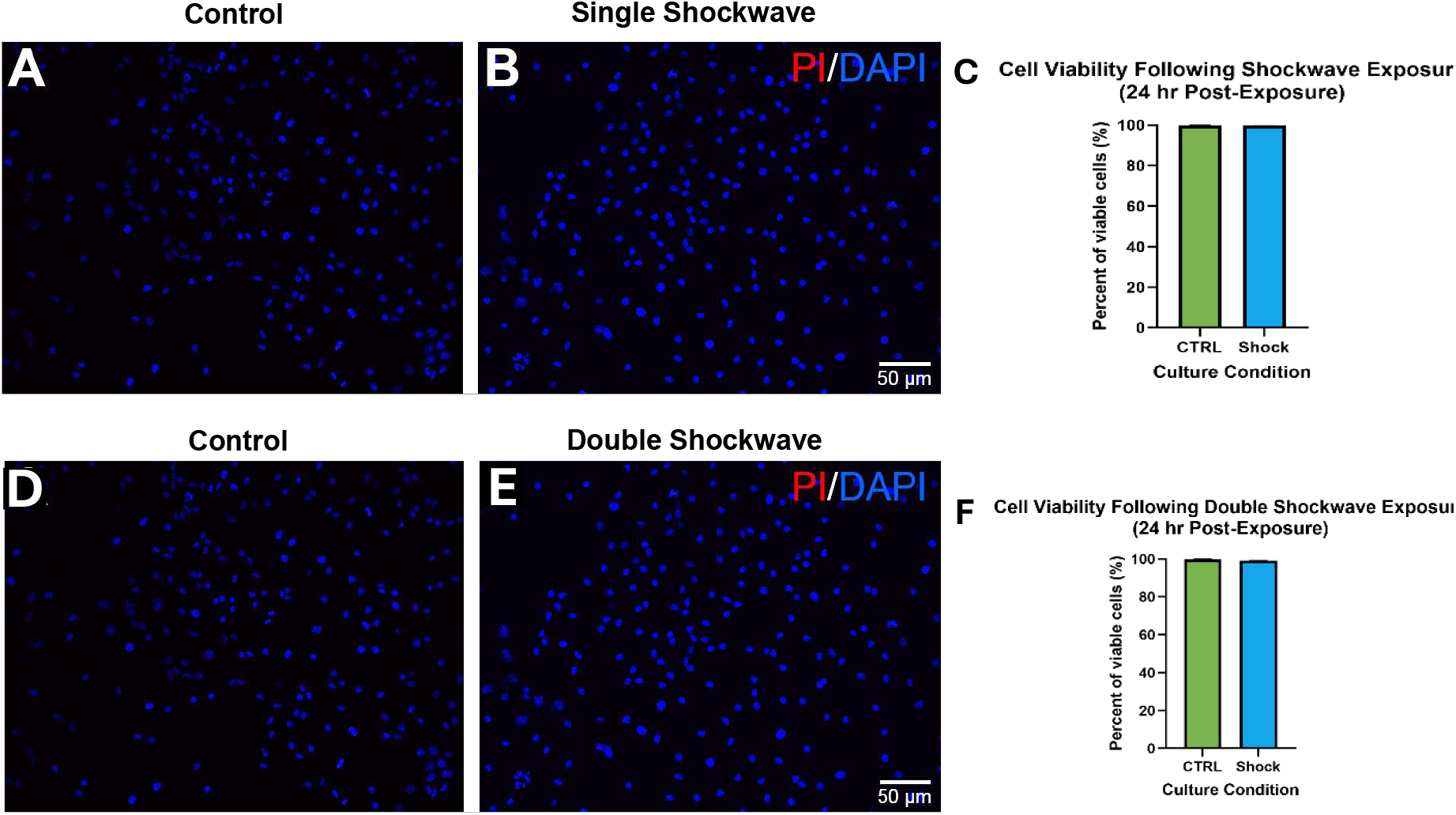
Comparison of Viability of AHPCs at 24 hr. Post-Exposure Following Single Shockwave Exposure from Top-to-Bottom. A-B: Representative fluorescence images of AHPC viability after 24 hr. post-exposure following single shockwave exposure. C: At 24 hrs. post-shockwave there was no significant difference between the culture conditions. N=2-3 independent experiments, 20-30 image fields were quantified for each condition. **Comparison of Viability of AHPC at 24 hr. Post-Exposure Following Double Shockwave Exposure from Above**. D-E: Representative fluorescence images of AHPC viability after 24 hr. post-exposure following double shockwave exposure. F: At 24 hours post-shockwave there was no significant difference between the culture conditions. N=3 independent experiments, 30 image fields were quantified for each condition. Bars represent the mean percentage of viable cells, and the error bars represent the standard error of the mean (±SEM).

**Fig. A3.**
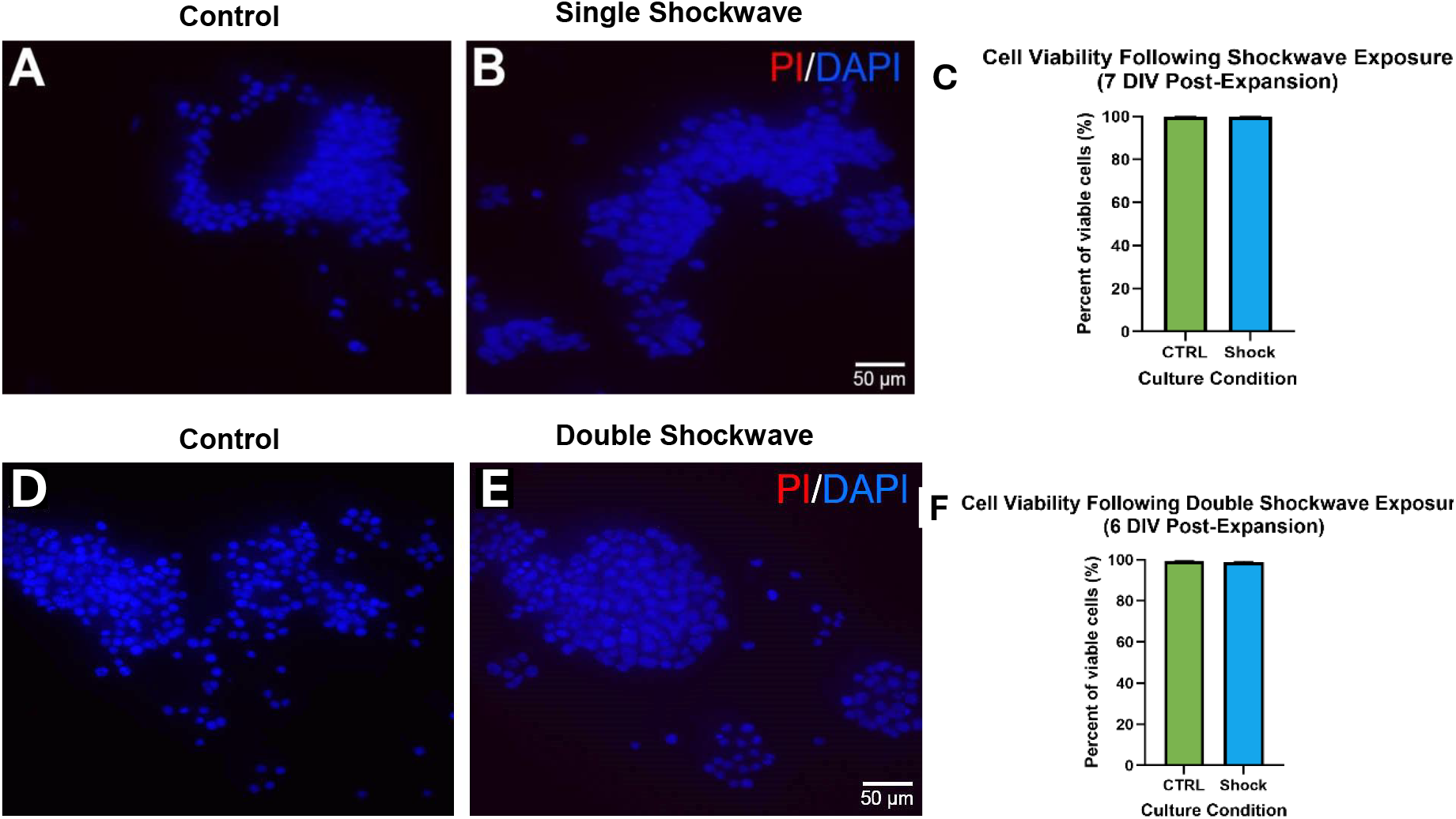
Comparison of Viability of AHPCs at 7 DIV Post-Expansion Following Single Shockwave Exposure from Top-to-Bottom. A-B: Representative fluorescence images of AHPC viability after 7 DIV post-expansion following single shockwave exposure. C: There was no significant difference between the culture conditions. N=2-3 independent experiments, 20-30 image fields were quantified for each condition. **Comparison of Viability of AHPCs at 6 DIV Post-Expansion Following Double Shockwave Exposure from Top-to-Bottom**. D-E: Representative fluorescence images of AHPC viability after 6 DIV post-expansion following single shockwave exposure. F: There was no significant difference between the culture conditions. N=3 independent experiments, 30 image fields were quantified for each condition. Bars represent the mean percentage of viable cells, and the error bars represent the standard error of the mean (±SEM).

**Fig. A4.**
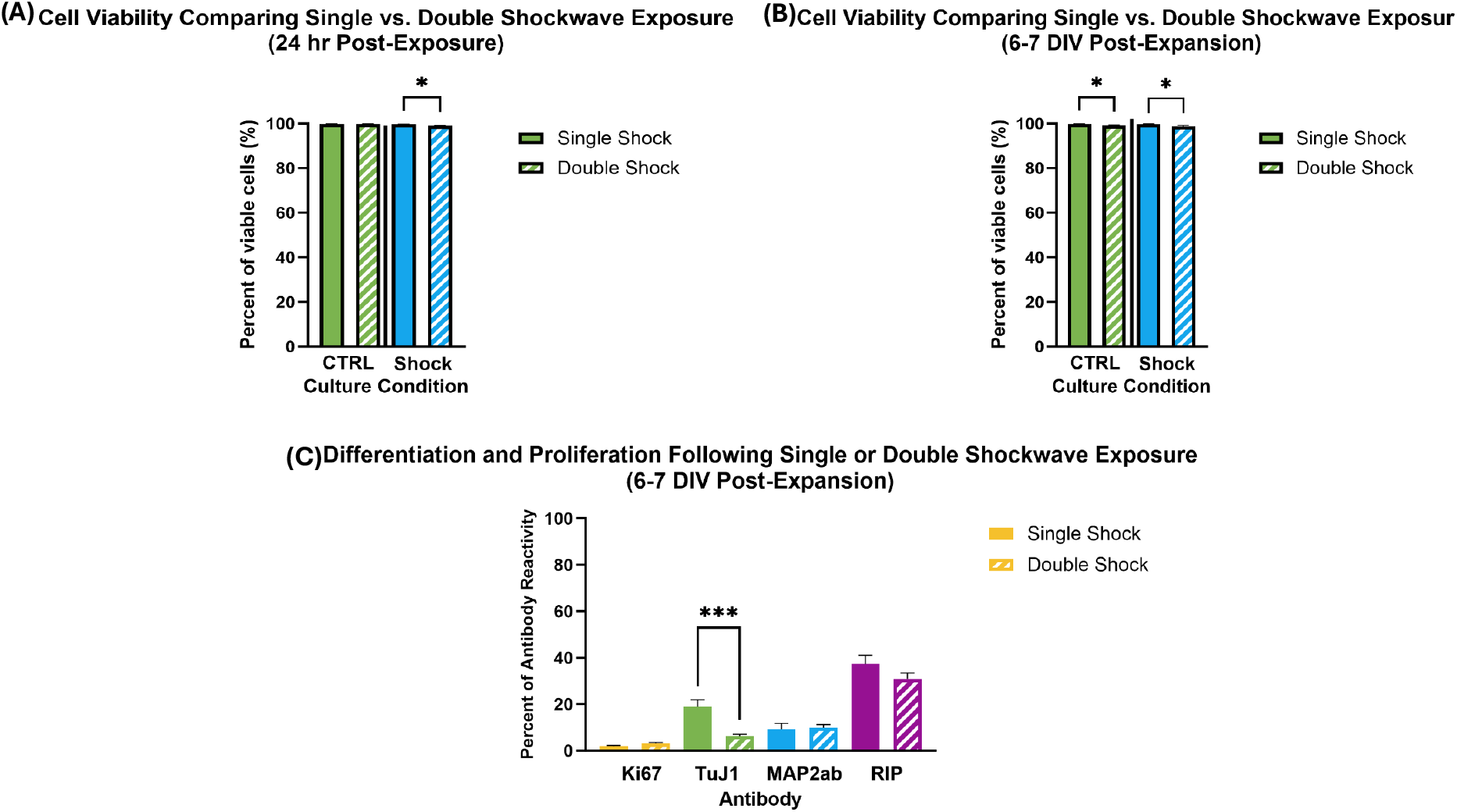
(A) Comparison of Viability of AHPCs at 24-hrs Post-Exposure Following Single or Double Shockwave Exposure from Top to Bottom. Quantitative analysis of AHPCs at 24-hrs. post-exposure following single or double shockwave exposure from above. There was a significant decrease in cell viability following double shockwave exposure compared to single shockwave exposure. There was no significant difference between the control culture conditions. N=3 independent experiments, 30 image fields were quantified for each condition. **(B) Comparison of Viability of AHPCs at 6-7 DIV Post-Expansion Following Single or Double Shockwave Exposure from Top to Bottom**. Quantitative analysis of AHPCs at 6-7 DIV post-expansion following single or double shockwave exposure from above. There was a significant decrease in cell viability following double shockwave exposure compared to single shockwave exposure for both the control and exposed culture conditions. N=3 independent experiments, 30 image fields were quantified for each condition. **(C) Comparison of Differentiation and Proliferation of AHPCs at 4 DIV Post-Expansion Following Single or Double Shockwave Exposure from Top to Bottom**. Quantitative analysis of AHPCs at 6-7 DIV post-expansion following single or double shockwave exposure from above. There was a significant decrease in the expression of TuJ1 following double shockwave exposure compared to single shockwave exposure. There was no significant difference between exposure conditions for the other antibodies tested. N=2-3 independent experiments, 20-30 image fields were quantified for each condition. Bars represent the mean percentage of viable cells, and the error bars represent the standard error of the mean (±SEM).

**Fig. A5.**
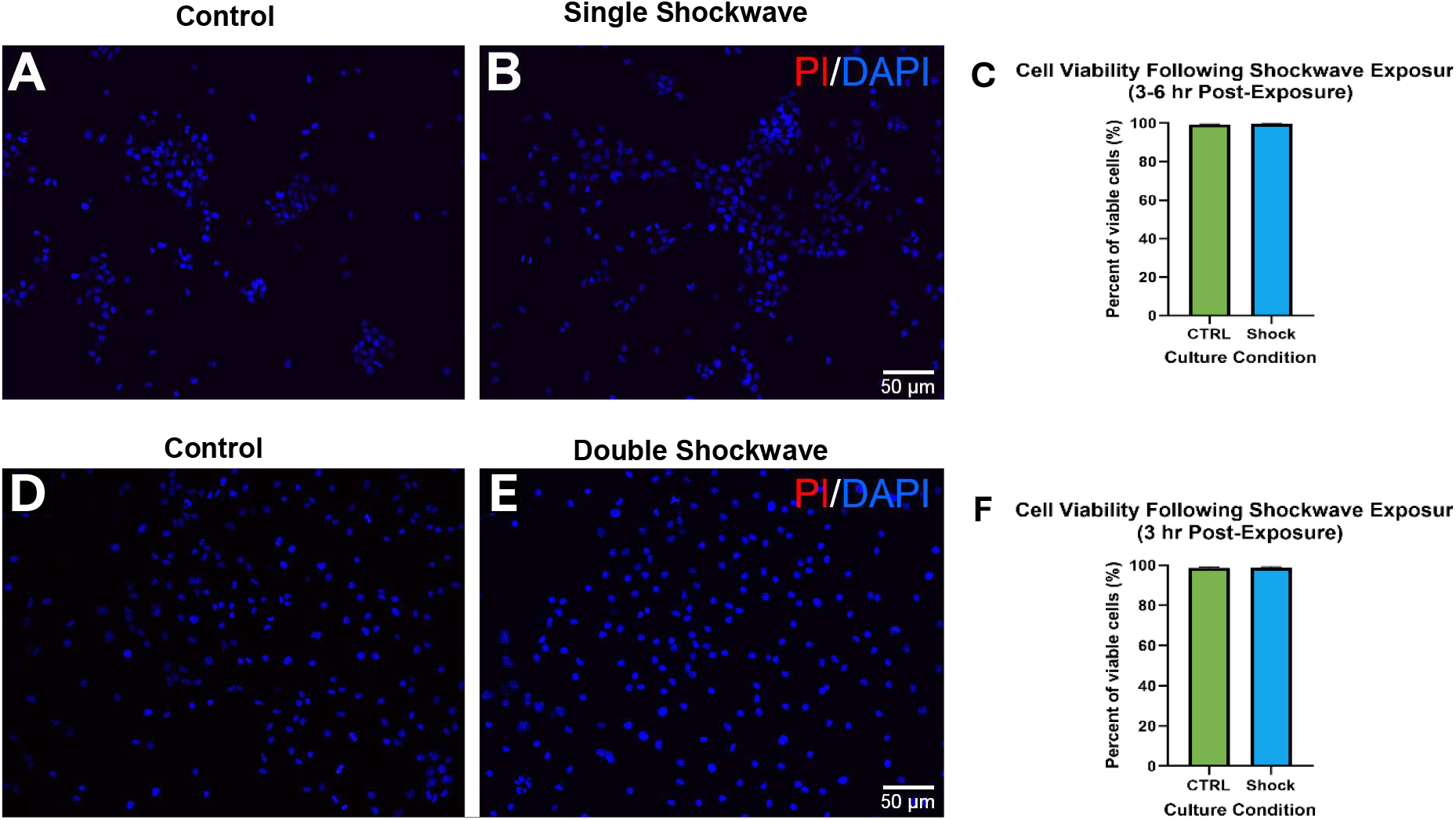
Comparison of Viability of AHPCs at 3 hr. Post-Exposure Following Single Shockwave Exposure from Bottom-to-Top. A-B: Representative fluorescence images of AHPC viability after 3 hr. post-exposure following single shockwave exposure. C: At 24 hours post-shockwave there was no significant difference between the culture conditions. N=3 independent experiments, 30 image fields were quantified for each condition. **Comparison of Viability of AHPC at 3 hr. Post-Exposure Following Double Shockwave Exposure from Below**. D-E: Representative fluorescence images of AHPC viability after 3 hr. post-exposure following double shockwave exposure. F: At 24 hours post-shockwave, there was no significant difference between the culture conditions. N=3 independent experiments, 30 image fields were quantified for each condition. Bars represent the mean percentage of viable cells, and the error bars represent the standard error of the mean (±SEM).

**Fig. A6.**
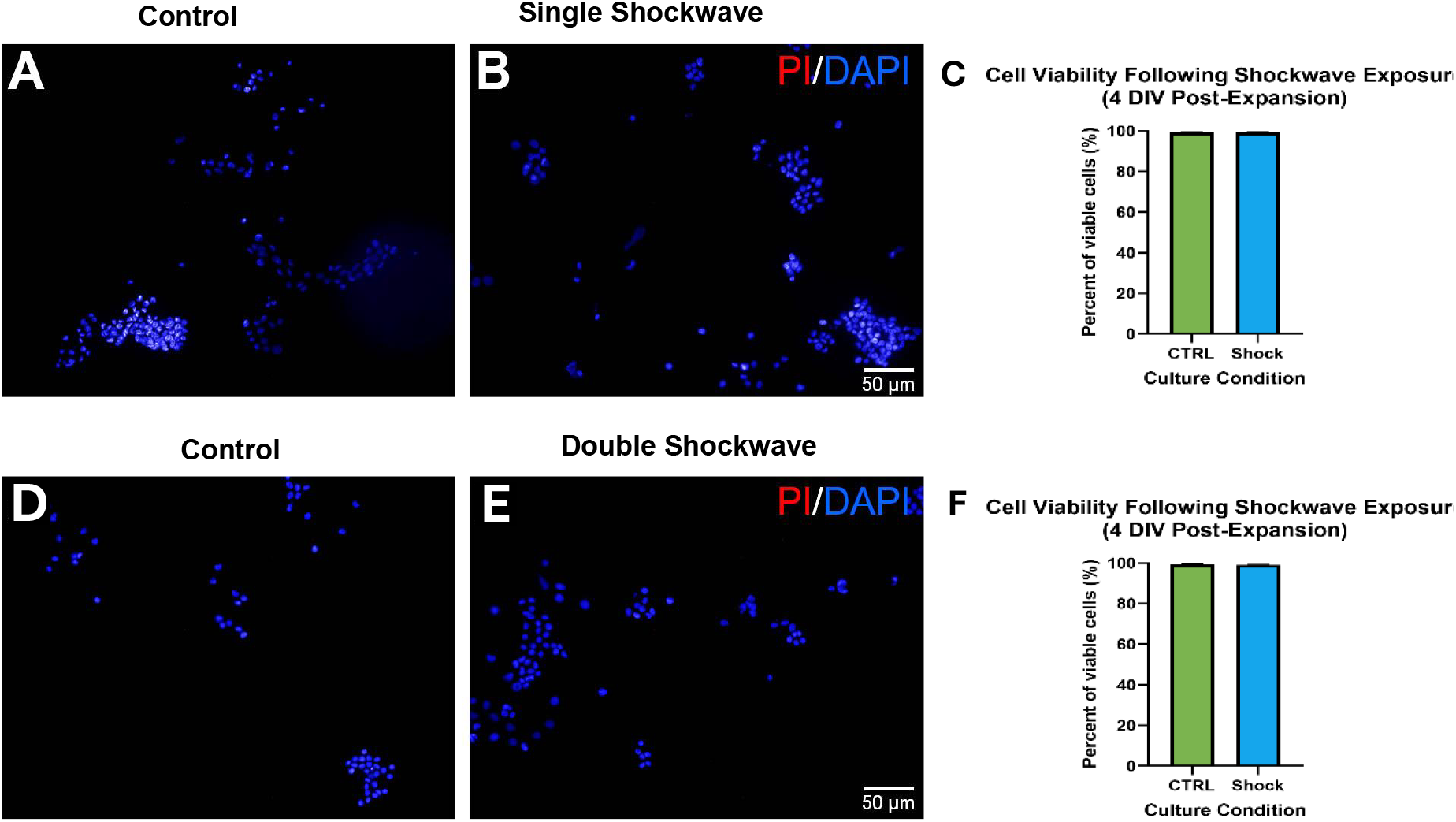
Comparison of Viability of AHPCs at 4 DIV Post-Expansion Following Single Shockwave Exposure from Bottom to Top. A-B: Representative fluorescence images of AHPC viability after 4 DIV post-expansion following single shockwave exposure. C: There was no significant difference between the culture conditions. **Comparison of Viability of AHPCs at 4 DIV Post-Expansion Following Double Shockwave Exposure from Bottom to Top**. D-E: Representative fluorescence images of AHPC viability after 4 DIV post-expansion following single shockwave exposure. F: There was no significant difference between the culture conditions. Bars represent the mean percentage of viable cells, and the error bars represent the standard error of the mean (±SEM). N=3 independent experiments, 30 image fields were quantified for each condition.

**Fig. A7.**
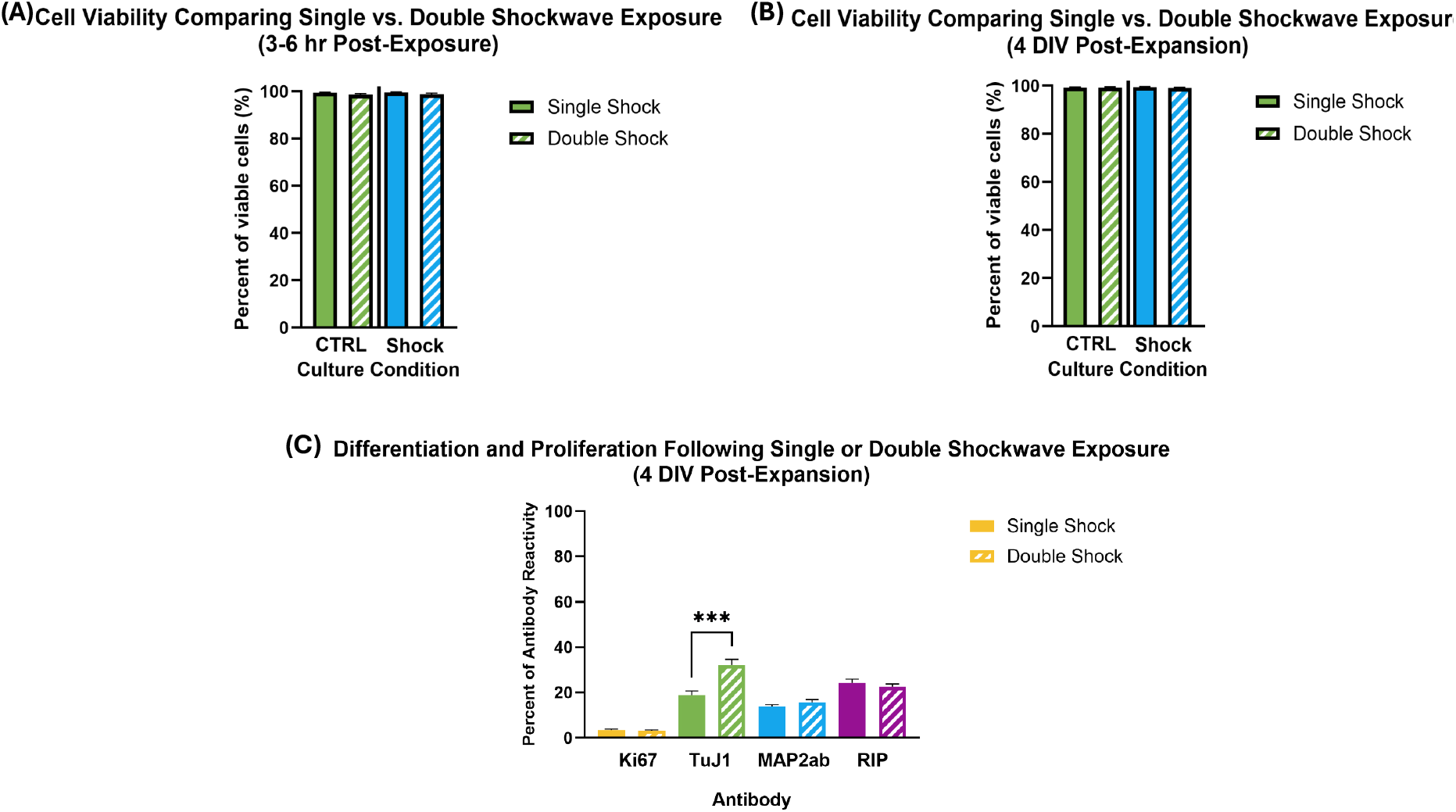
(A)Comparison of Viability of AHPCs at 3-6-hrs. Post-Exposure Following Single or Double Shockwave Exposure from Bottom to Top. Quantitative analysis of AHPCs at 3-6-hrs. post-exposure following single or double shockwave exposure from below. There was no significant difference between the control or shockwave culture conditions for either exposure condition. **(B)Comparison of Viability of AHPCs at 4 DIV Post-Expansion Following Single or Double Shockwave Exposure from Bottom to Top**. Quantitative analysis of AHPCs at 4 DIV post-expansion following single or double shockwave exposure from below. There was no significant difference between the control or shockwave culture conditions for either exposure condition. (C) **Comparison of Differentiation and Proliferation of AHPCs at 4 DIV Post-Expansion Following Single or Double Shockwave Exposure from Bottom to Top**. Quantitative analysis of AHPCs at 6-7 DIV post-expansion following single or double shockwave exposure from below. There was a significant increase in the expression of TuJ1 following double shockwave exposure compared to single shockwave exposure. There was no significant difference between exposure conditions for the other antibodies tested. N=3 independent experiments, 30 image fields were quantified for each conditions in plots. Bars represent the mean percentage of viable cells, and the error bars represent the standard error of the mean (±SEM).

**Fig. A8.**
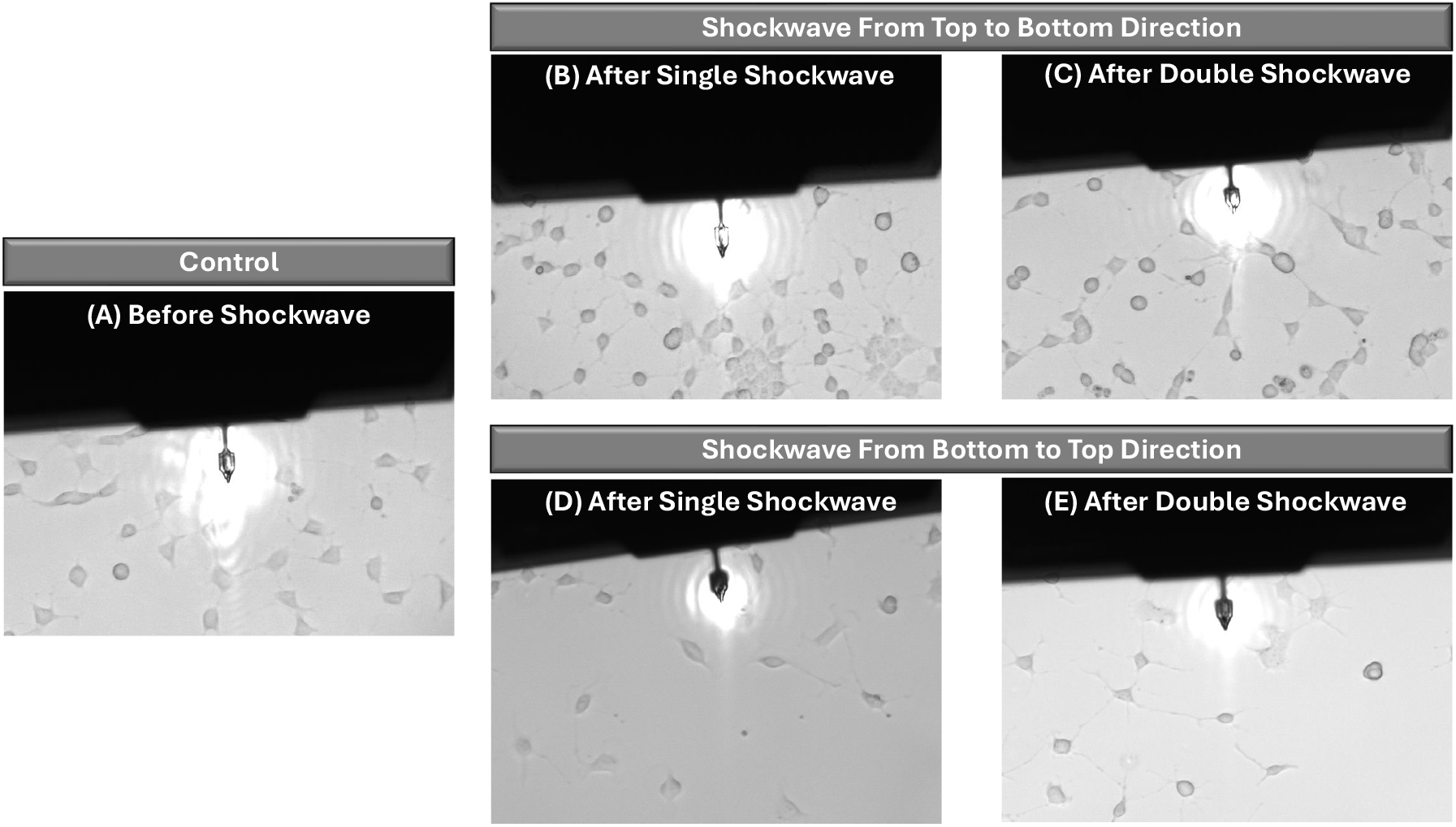
Phase contrast images of AHPC control cells and cells exposed to shockwave, both kept in the incubator for 3 hours, showed the restoration process. By observing these images, cells were recovering and their cytoskeletal structures and gradual return to regular cell morphology.

